# The auditory midbrain mediates tactile vibration sensing

**DOI:** 10.1101/2024.03.08.584077

**Authors:** Erica L. Huey, Josef Turecek, Michelle M. Delisle, Ofer Mazor, Gabriel E. Romero, Malvika Dua, Zoe K. Sarafis, Alexis Hobble, Kevin T. Booth, Lisa V. Goodrich, David P. Corey, David D. Ginty

## Abstract

Vibrations are ubiquitous in nature, shaping behavior across the animal kingdom. For mammals, mechanical vibrations acting on the body are detected by mechanoreceptors of the skin and deep tissues and processed by the somatosensory system, while sound waves traveling through air are captured by the cochlea and encoded in the auditory system. Here, we report that mechanical vibrations detected by the body’s Pacinian corpuscle neurons, which are unique in their ability to entrain to high frequency (40-1000 Hz) environmental vibrations, are prominently encoded by neurons in the lateral cortex of the inferior colliculus (LCIC) of the midbrain. Remarkably, most LCIC neurons receive convergent Pacinian and auditory input and respond more strongly to coincident tactile-auditory stimulation than to either modality alone. Moreover, the LCIC is required for behavioral responses to high frequency mechanical vibrations. Thus, environmental vibrations captured by Pacinian corpuscles are encoded in the auditory midbrain to mediate behavior.

## Introduction

Vibration sensing is ubiquitous across the animal kingdom, enabling organisms to interpret their dynamic surroundings, communicate, detect predators and prey, and navigate their environment^1–6^. In mammals, vibrations that propagate through the air in the form of sound waves are transduced into electrical signals in the cochlea and encoded by the auditory system^7^. Analogously, mechanical vibrations acting on the body are detected by fast conducting low-threshold mechanoreceptors (Aβ LTMRs) of the body and encoded by the somatosensory system^8,9^. In many animals, including humans, the vibration frequency detection range of the auditory system (20 Hz-20 kHz) partially overlaps with that of the somatosensory system (1-1000 Hz)^8,10–12^. In some smaller mammals, such as mice, the encoding of vibration frequency in the somatosensory system (1-1000 Hz) and auditory system (∼1 khz-100 kHz) is largely non-overlapping^10,11,13–16^. The prevailing view is that vibration encoding in the ascending somatosensory and auditory pathways is anatomically and functionally distinct.

Our understanding of LTMRs and their role in detecting mechanical vibrations mainly arises from experiments in which vibratory stimuli are presented directly to the body; however, the somatosensory system can also detect subtle, low amplitude environmental vibrations. In a noteworthy example, elephants can communicate over miles, sensing vocalizations and movements of conspecifics by detecting vibrations of the ground using mechanoreceptors in their feet and trunks^4,17^. Among the LTMRs, Pacinian corpuscle Aβ rapidly adapting type-2 (RA2)-LTMRs are unique in their capacity to encode high frequency vibrations^8^. In humans and primates, Pacinians are located in the deep dermis of glabrous skin and also associated with joints and connective tissue^18,19^. In rodents, Pacinians are enriched in the periosteum of wrist and ankle bones, rendering them responsive to vibrations transmitted through the skeleton^8,14,15,20,21^. Recordings from Pacinian afferents in awake, behaving mice have recently revealed their exquisite sensitivity to self-movements and substrate vibrations under naturalistic conditions^15^. Indeed, Pacinian corpuscle Aβ RA2-LTMRs can fire robustly in response to low-amplitude environmental vibrations, including those initiated by gentle movements on a surface located meters from where an animal stands^15,22–25^.

The canonical view is that the neural impulses originating in the periphery and encoding mechanical vibrations acting of the body are conveyed via the brainstem dorsal column nuclei (DCN) to the ventroposterolateral thalamus (VPL) *en route* to primary somatosensory cortex (S1)^9,26^. However, our recent work suggests that mechanical vibration sensitive DCN neurons project to the inferior colliculus^27^, a midbrain region involved in auditory information processing^7^. The inferior colliculus (IC) is comprised of multiple subregions: the central nucleus of the inferior colliculus (CNIC), which receives auditory input in a tonotopic manner, and the lateral cortex of the inferior colliculus (LCIC), which receives inputs from multiple sensory modalities, including somatosensory inputs from the DCN (Figure 1A, Figure S1)^7,27–38^. Here, we use electrophysiological recordings, genetic manipulations, and behavioral analyses to show that most LCIC neurons of the auditory midbrain encode both high frequency mechanical vibrations captured by Pacinian corpuscle LTMRs and auditory vibrations captured by the cochlea to mediate behavioral responses to environmental vibrations.

**Figure 1.**
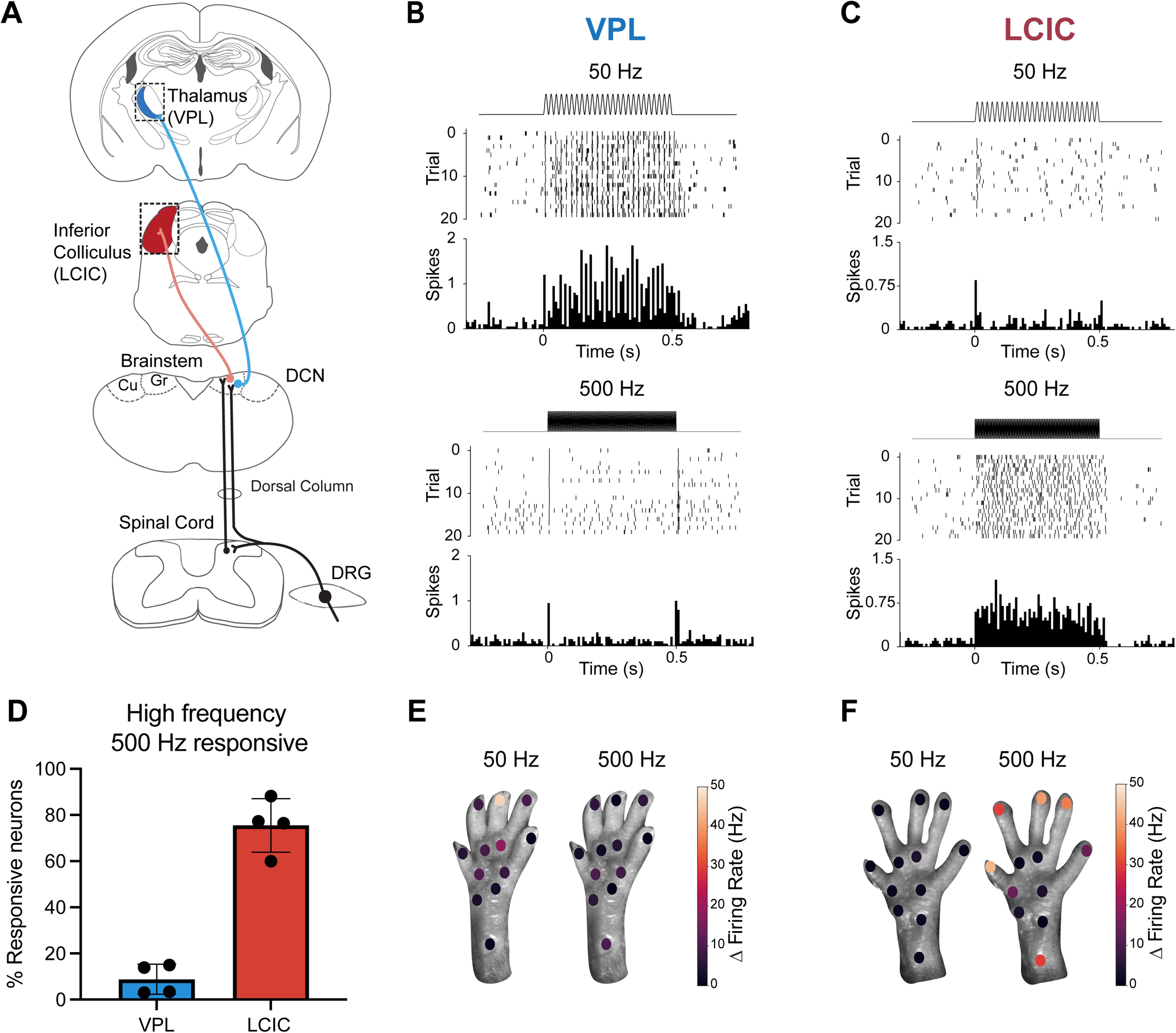
LCIC and VPL neurons are differentially tuned to mechanical vibratory stimuli. **A)** The brainstem dorsal column nuclei (DCN) receive inputs from dorsal root ganglia (DRG) primary sensory neurons and spinal cord projection neurons. Non-overlapping subsets of DCN projection neurons target the ventroposterolateral thalamus (VPL; blue) and the external nucleus of the inferior colliculus (LCIC; red). **B and C)** Representative VPL (B) and LCIC (C) neuron spike raster and histogram in response to 50 Hz and 500 Hz vibratory stimuli delivered at a 10 mN force to the hindlimb. Top: spike raster; bottom: average histogram in 10 ms bins. **E)** Percentage of neurons (mean ± SD) with hindbody brush receptive fields (tested by brushing the hindpaw, thigh, and trunk) that respond to 500 Hz stimulation of the limb at a 10 mN force in the VPL (n= 182 neurons from 4 animals) and LCIC (n= 76 neurons from 4 animals). Each data point corresponds to measurements in one animal. See also Supplementary Figure 3. **E and F)** Example VPL (E) and LCIC (F) neuron receptive field organization for 50 Hz and 500 Hz stimulation to multiple regions across the hindlimb delivered at a 10 mN intensity. Stimulus locations are superimposed on images of the hindpaw.

## Results

### LCIC and VPL neurons are differentially tuned to mechanical vibrations

Distinct populations of DCN neurons send axonal projections to either the VPL of the thalamus or the LCIC of the inferior colliculus^37^. DCN axonal terminals from the gracile nucleus (representing the hindbody) and the cuneate nucleus (representing the forebody) segregate to different regions of the VPL but innervate overlapping regions within the LCIC (Figure S1). To compare how mechanical vibrations acting on the body are represented in the LCIC and VPL, we performed *in vivo* multi-electrode array (MEA) recordings in these brain regions in urethane anesthetized animals (Figure S2A,E). We first brushed the animal to identify neurons containing limb receptive fields. Sinusoidal mechanical vibrations of different frequencies and amplitudes were then delivered to the limbs using a mechanical stimulator. While the LCIC and VPL both contain forelimb-and hindlimb-responsive neurons (Figure 1B-C, Figure S2), we focused primarily on hindlimb responsive neurons. Vibrations were delivered to the most responsive region, or “hotspot,” of a neuron’s receptive field. In the VPL, vibratory receptive fields tended to be focal areas such as an individual pad or digit (Figure 1E). In contrast, receptive fields of LCIC neurons lacked a focal hotspot and were spatially extensive (Figure 1F). In these cases, the mechanical stimulator was positioned on the heel of the hindlimb. Vibratory stimuli were then delivered at frequencies between 10-900 Hz and forces between 2-50 mN. The majority of VPL neurons responded robustly to low frequency vibrations (<200 Hz), but only transiently to high frequency vibrations (>200 Hz; Figure 1B, Figure S2C-D). While their pre-synaptic DCN inputs are able to entrain (phase-lock firing to each cycle of the stimulus) to vibratory frequencies up to 100 Hz^27^, no VPL neurons could entrain to vibratory frequencies up to 100 Hz, although some neurons could entrain up to 50 Hz. Strikingly, LCIC neurons showed an inverse response pattern compared to VPL neurons, with most neurons responding transiently to low frequency vibrations (<200 Hz) but firing robustly to high frequency stimuli (200-900 Hz; Figure 1C, Figure S2G-H, Figure S5I,M). While many DCN projection neurons targeting the LCIC entrained to vibratory frequencies from 10-500 Hz^27^, LCIC neurons only transiently responded to low frequency stimuli (Figure 1C, Figure S2G-H), suggesting a synaptic filtering mechanism between the DCN and LCIC that preferentially favors high frequency vibratory stimuli. Furthermore, although LCIC neurons responded robustly to high frequency vibrations, they exhibited little or no entrainment (Figure 1C, Figure S2G-H).

While high frequency vibratory responses were abundant throughout the LCIC, the sparsity of high frequency responses detected in the VPL was surprising and could be attributed to missing a potential “high frequency hotspot” region in the VPL in our recordings. Therefore, considering that the VPL is an elongated structure extending along the anterior-posterior axis of the brain, we next used a four shanked MEA to perform a detailed mapping of somatotopy across the VPL to search for high frequency vibratory responses. We recorded from a minimum of 9 penetrations per animal using the four-shanked probe, enabling sampling across at least 36 regions of the VPL, distributed across the medio-lateral and anterior-posterior axes. The resulting somatotopic map of the VPL aligned well with those previously reported, with receptive fields moving from forelimb to hindlimb to trunk, when moving from medial to lateral within the VPL (Figure S3A,C-D)^39–42^. A larger proportion of VPL neurons responded to brush of the hindpaw compared to the thigh or the trunk (Figure S3F). This emphasis on hindpaw representation was less pronounced in LCIC neurons, which often exhibited expansive receptive fields spanning multiple body regions (Figure S3B, G). For VPL neurons with brush responses to hindbody regions, high frequency vibration of the hindlimb activated ∼9 percent of neurons, and these high frequency responsive neurons were sparsely distributed throughout the VPL, not concentrated in a particular subregion (Figure 1D, Figure S3E). In contrast, >75 percent of brush responsive LCIC neurons responded robustly to high frequency vibrations, indicating that high frequency vibratory information is particularly salient in the LCIC (Figure 1D).

To assess the relationship between mechanical force thresholds and vibration frequency in the VPL and LCIC, we next obtained force-frequency tuning curves for vibration sensitive VPL and LCIC neurons by determining force thresholds for individual neurons across a range of vibratory frequencies. The force threshold for a given frequency was defined as the minimum force required to elicit 20 percent of a neuron’s maximum firing rate. The tuning curves for VPL and LCIC neurons were approximately U-shaped, although the most sensitive regions of their respective tuning curves were non-overlapping: VPL neurons had the lowest force thresholds to vibrations between 50-200 Hz and LCIC neurons had the lowest force thresholds to vibrations between 300-500 Hz (Figure 2A-E). Thus, most LCIC neurons are activated by high frequency vibratory stimuli and exhibit expansive receptive fields, while most VPL neurons are preferentially activated by low frequency stimuli in a spatially restricted manner.

**Figure 2.**
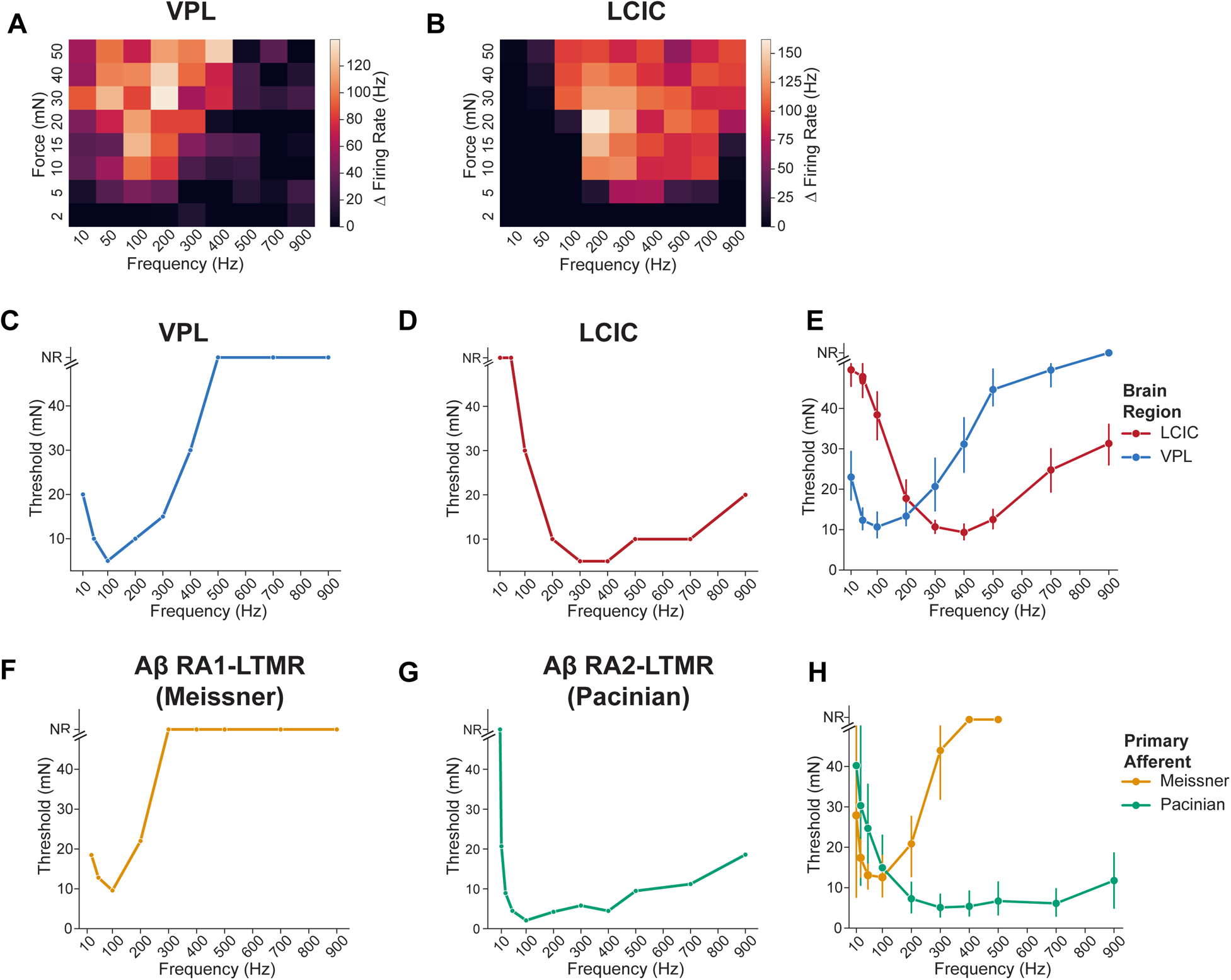
LCIC and VPL neurons exhibit distinct force-frequency tuning curves. **A and B)** Representative VPL (A) and LCIC (B) neuron responses to vibrational frequencies delivered at different forces to the hindlimb. Heatmap intensity corresponds to the neuron’s average change in firing to the sustained portion of vibratory stimuli (200 ms window beginning 50 ms after stimulus onset). **C and D)** Force-frequency threshold curves obtained from heatmaps in A and B, where the force threshold for a vibrational frequency is defined as the minimum force required to elicit 20% of the neuron’s maximum firing during the sustained portion of the stimulus. Frequencies in which the highest force tested (50 mN) failed to reach threshold are designated NR (non-responsive). **E)** Average threshold curves (mean ± 95% confidence interval) for neurons responsive to hindlimb vibration recorded in the VPL (n=15 neurons from 6 animals) and LCIC (n=22 neurons from 6 animals). **F and G)** Force-frequency threshold curves for representative Meissner corpuscle innervating Aβ RA1-LTMR afferent (F) and Pacinian corpuscle innervating Aβ RA2-LTMR afferent (G). Threshold is defined as the minimum force necessary for the neuron to reach 50% vibratory entrainment. Frequencies in which the highest force tested (50 mN) failed to reach threshold are designated NR (non-responsive). **H)** Average threshold curves (mean ± 95% confidence interval) for hindlimb Meissner Aβ RA1-LTMRs (n=4 neurons from 4 animals) and Pacinian Aβ RA2-LMTRs (n=6 neurons from 5 animals).

### LCIC vibration responses are mediated by Pacinian corpuscle Aβ RA2-LTMRs

The distinct frequency tuning curves and receptive field properties of VPL and LCIC neurons prompted us to ask whether these brain regions receive inputs from different primary mechanosensory neuron types. Rapidly adapting Aβ LTMRs exhibit distinct force-frequency tuning relationships and receptive fields. Most rapidly adapting LTMRs are optimally sensitive to low frequency vibratory stimuli (40-150 Hz) and exhibit relatively small receptive fields^8,9,13,43–45^ This includes Meissner corpuscle Aβ RA1-LTMRs, enriched in glabrous skin of pedal pads and digits, hair follicle Aβ RA-LTMRs associated with body hairs, and Krause corpuscle Aβ RA-LTMRs of the genitalia^8,13,44,46^. In contrast, Pacinian corpuscle Aβ RA2-LTMRs, which in mice are enriched in the periosteum of ankle and wrist bones, are most sensitive to high frequency vibratory stimuli (50-1000 Hz) and have extremely large receptive fields^15,20,47–52^. To directly compare the tuning of Meissner corpuscle Aβ RA1-LTMRs and Pacinian corpuscle Aβ RA2-LTMRs to VPL and LCIC neurons, we performed *in vivo* extracellular recordings of DRG neurons and measured force-frequency tuning curves of Meissner and Pacinian afferents (Figure 2F-H). As predicted, the force-frequency tuning curves of Pacinian corpuscle Aβ RA2-LTMRs resembled those of LCIC neurons, while the tuning curves of Meissner corpuscle Aβ RA1-LTMRs more closely resembled VPL neurons (Figure 2F-H). These findings led us to hypothesize that Pacinian afferents have an outsized role in shaping the response properties of most LCIC neurons, while Meissner corpuscle afferents and other Aβ RA-LTMRs of the body preferentially contribute to the response properties of most VPL neurons.

To directly test the relationship between primary sensory neuron activity and vibration tuning in the LCIC and VPL, we utilized mouse genetic tools to assess the contributions of Pacinian corpuscles and Meissner corpuscles to LCIC and VPL responses. Ret signaling in somatosensory neurons is necessary for formation of Pacinian corpuscles, and thus *Avil^Cre^; Ret^floxGFP/floxGFP^* (Ret cKO) mice, which lack *Ret* in DRG neurons, lack Pacinian corpuscles (Figure 3B, Figure S4F)^53,54^. Conversely, TrkB signaling is necessary for formation of Meissner corpuscles, and thus *Avil^Cre^; TrkB^flox/flox^* (TrkB cKO) mice lack Meissner corpuscles (Figure 3A)^13,44,55^. Importantly, Ret cKO mice have a normal density of Meissner corpuscles and TrkB cKO mice have a normal density of Pacinian corpuscles (Figure S4).

**Figure 3.**
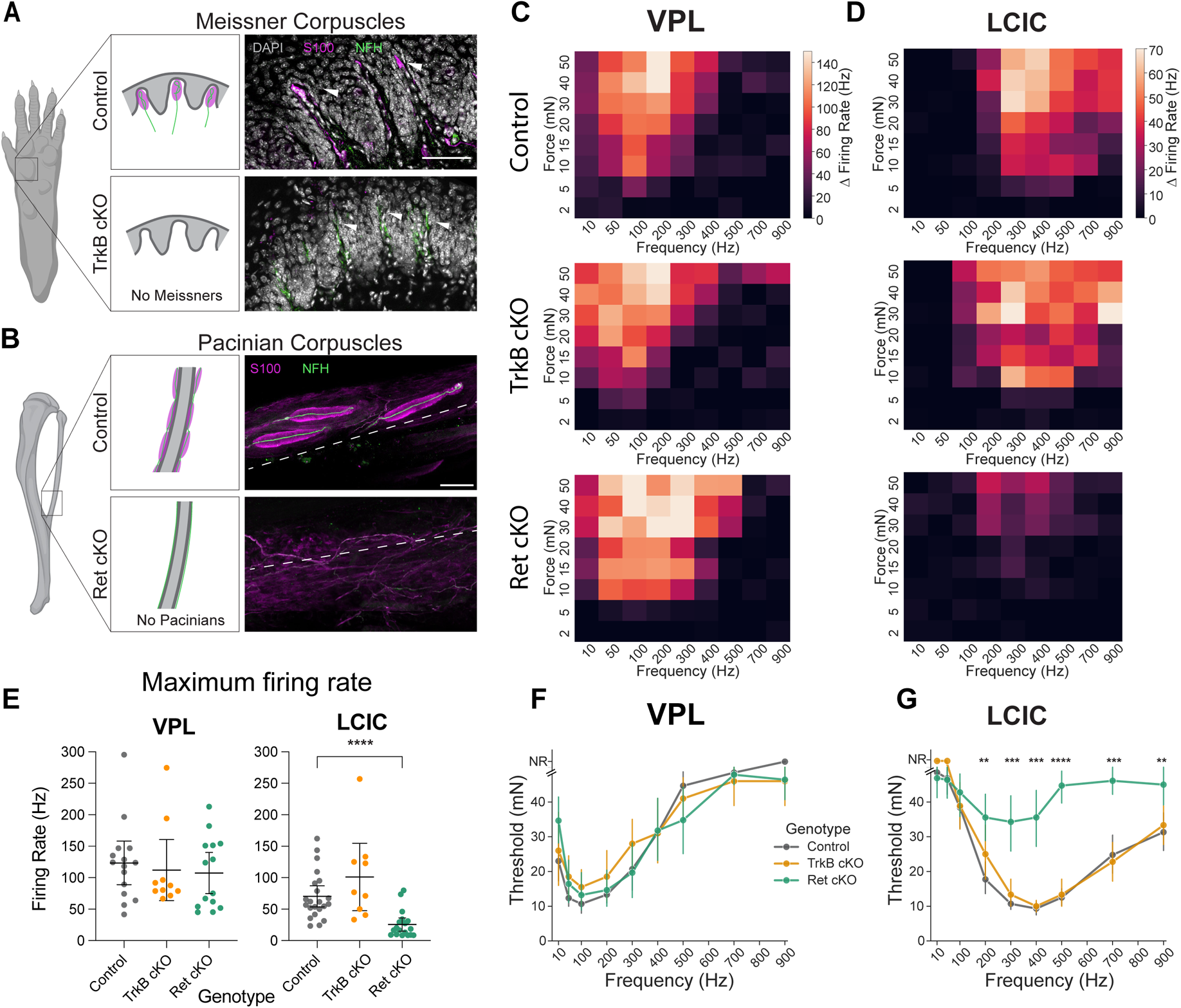
LCIC vibration responses are mediated by Pacinian corpuscle Aβ RA2-LTMRs. **A)** Meissner corpuscles, located in the plantar pad and digit glabrous skin, are absent in *Avil^Cre^; TrkB^flox/flox^*(TrkB cKO) mice, which lack TrkB in DRG sensory neurons. Images of skin histology from a TrkB cKO and littermate control, stained for anti-NFH (labels myelinated axons, green) and anti-S100 (labels glial cells, magenta). Scale bar = 50 µm. See also Supplementary Figure 4 for quantification. **B)** Pacinian corpuscles, located in the periosteum of the fibula, are absent in *Avil^Cre^; Ret^flox/flox^* (Ret cKO) mice, which lack Ret in DRG sensory neurons. Images of whole mount histology of the limb of a Ret cKO animal and littermate control, stained for anti-NFH and anti-S100 to visualize Pacinian corpuscles. White dashed line indicates position of the fibula. Scale bar = 100 µm. See also Supplementary Figure 4 for quantification. **C and D)** Force-frequency heat maps for representative VPL (C) and LCIC (D) neurons in control, TrkB cKO and Ret cKO animals. Heatmap intensity corresponds to the neuron’s average change in firing to the sustained portion of vibratory stimuli. **E)** Maximum firing rate for VPL and LCIC (mean ± 95% confidence interval) neurons to vibratory stimuli in control, TrkB cKO and Ret cKO mice. Firing rates of LCIC neurons in Ret cKO and control mice were significantly different (****p<0.0001 Mann-Whitney U test). Firing rates of VPL neurons in TrkB cKO, Ret cKO, and controls were not significantly different. Firing rates of LCIC neurons in TrkB cKO and controls were not significantly different (Mann-Whitney U test). **F)** Average threshold curves of VPL neurons (mean ± 95% confidence interval) recorded in control (n=15 neurons from 6 animals), TrkB cKO (n=10 neurons from 3 animals), and Ret cKO (n=14 neurons from 4 animals) mice. Thresholds between Ret cKO, TrkB cKO and controls were not significantly different (Mann-Whitney U Test with Bonferroni correction for multiple comparisons). **G)** Average threshold curves of LCIC neurons (mean ± 95% confidence interval) recorded in control (n=22 neurons from 6 animals), TrkB cKO (n=9 neurons from 4 animals), and Ret cKO (n=18 neurons from 4 animals) mice. (*p<0.05, **p<0.01, ***p<0.001, ****p<0.0001, Mann-Whitney U test with Bonferroni correction for multiple comparisons). While thresholds between Ret cKO and controls were significantly different, thresholds between TrkB cKO and controls were not (Mann-Whitney U Test with Bonferroni correction for multiple comparisons).

Surprisingly, VPL neurons recorded in TrkB cKO and Ret cKO mice showed no notable shifts in their vibration tuning curves (Figure 3C, F). While VPL neurons of TrkB cKO and Ret cKO mice exhibited a decrease in the magnitude of the offset response at certain vibration frequencies, the onset and sustained phases of the response revealed no significant differences (Figure S5A-H). Meissner corpuscle and Pacinian corpuscle afferents are not the only primary sensory neurons responsive to vibrations, and indeed Aβ RA-LTMRs associated with hair follicles and other end organs and Aβ SA1-LTMRs found in both hairy and glabrous skin also entrain to vibratory stimuli^8,52,56,57^. Therefore, it is likely that VPL neurons are sensitive to mechanical vibrations via inputs from both Meissner and Pacinian corpuscle LTMRs or other vibration sensitive LTMRs^55,58^. In stark contrast, vibration responsiveness of LCIC neurons was nearly absent in animals lacking Pacinian corpuscles, although LCIC responses were normal in mice lacking Meissner corpuscles (Figure 3D, E, G, Figure S5I-P). Taken together, these findings indicate that high frequency mechanical vibrations acting on the body and detected by Pacinian corpuscle Aβ RA2-LTMRs are prominently encoded in the LCIC.

### LCIC neurons integrate multimodal tactile-auditory stimuli in awake mice

Recent recordings from Pacinian afferents under awake conditions revealed that Pacinian Aβ RA2-LTMRs are exquisitely sensitive to environmental vibrations encountered during natural behaviors^15^. Because Pacinian corpuscles underlie vibration responses in LCIC neurons (Figure 3D, E, G) and our recent recordings in awake mice show that they are ultrasensitive to substrate vibrations^15^, we next asked whether LCIC neurons are also sensitive to low amplitude substrate vibrations in awake animals. We performed MEA recordings in awake head-fixed mice, but instead of applying vibration directly to the paw, we vibrated the platform the animal stood on (Figure 4A). Remarkably, in awake mice, 100 percent of LCIC neurons with hindlimb receptive fields responded robustly to 500 Hz stimulation of the surface platform, but few responded to a 50 Hz stimulus, similar to results from direct stimulation of the limb in recordings from anesthetized mice (Figure 4B-C). Thus, as with Pacinian corpuscle Aβ RA2-LTMRs, LCIC neurons encode low-amplitude, high frequency environmental vibrations.

**Figure 4.**
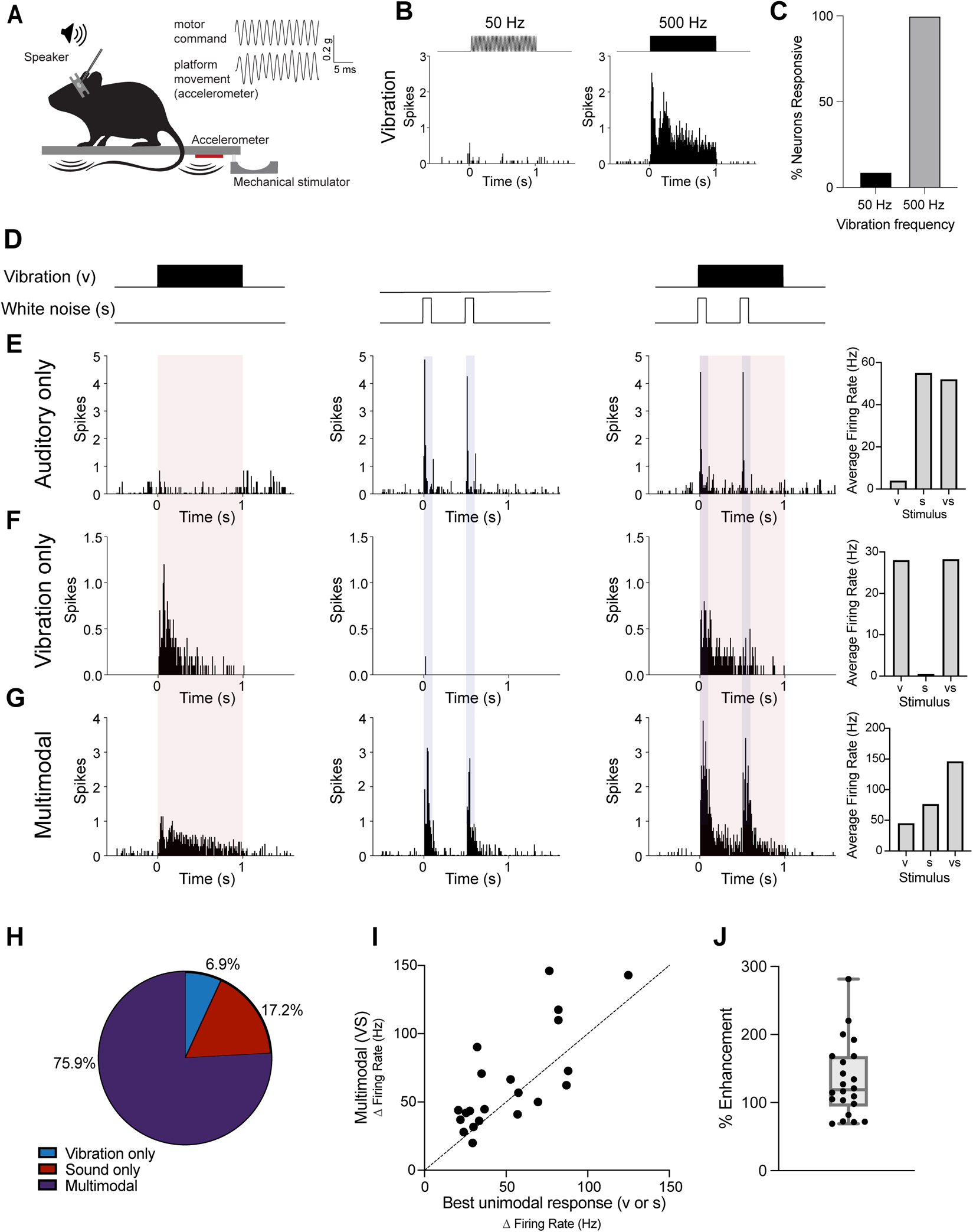
Audio-tactile multimodality of LCIC neurons in awake mice. **A)** Recordings of LCIC neurons were conducted in awake head-fixed mice as they stood on an acrylic platform, which could be stimulated at different vibratory frequencies. An accelerometer was attached underneath the acrylic platform to measure vibration frequency and amplitude. An example motor command and accelerometer signal are shown for 500 Hz vibration of the platform. A speaker is positioned on the contralateral side of the head from the recording site. **B)** Peristimulus time histogram (PSTH) for a representative LCIC neuron, aligned to 50 Hz (left) and 500 Hz (right) vibration stimulation of the acrylic platform at a vibration amplitude of ∼0.2 g (gravitational acceleration). PSTHs are baseline subtracted and reported in 10 ms bins. **C)** Percentage of hindlimb sensitive LCIC neurons that responded to 50 Hz and 500 Hz vibratory stimuli (n=24 neurons from 5 animals). **D)** 500 Hz substrate vibration (amplitude 0.2 g) and auditory white noise (frequency 0-50 kHz, 100 ms pulses, 65 dB SPL) were delivered in isolation and coincidentally to awake head-fixed mice. **E-G)** Example PSTHs from LCIC neurons that respond to auditory stimuli (E), vibration (F), or both (G), aligned to vibration (v), sound (s) and convergent (vs) stimuli. PSTHs are baseline subtracted and in 10 ms bins. Right: The example neuron’s average firing (during 0-0.1 s and 0.5-0.6 s windows) for each stimulus type. **H)** Percentage of neurons in control animals that respond to 500 Hz vibration only, sound only, or convergent stimuli (n=29 neurons from 5 animals). **I)** Scatter plot of firing rates in response to combined vibration and sound (vs) plotted against best unimodal response (vibration only or sound only). Unitary line denotes equal firing between best unimodal stimulus and combined stimulus conditions (n=21 neurons from 5 animals). **J)** Quantification of multisensory enhancement (corresponding to I), calculated using the following equation: 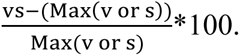. Neurons above 100% display multisensory enhancement (n=21 neurons from 5 animals)^76,77^.

The inferior colliculus is primarily an auditory nucleus, and the LCIC receives extensive auditory input from brainstem auditory nuclei^7^, prompting us to ask whether LCIC neurons responsive to high frequency tactile vibrations also respond to sound. Prior findings indicate that many LCIC neurons are broadly tuned to auditory stimuli, and most auditory-sensitive LCIC neurons respond to short pulses of broadband noise^30,35,59,60^. Therefore, we exposed animals to 100 ms pulses of a broadband white noise stimulus (0-50 kHz) at a 65 dB sound pressure level (SPL) with the goal of activating LCIC neurons that receive auditory inputs. We delivered 500 Hz mechanical vibrations to the platform surface and white noise pulses in isolation, as well as coincident tactile-auditory stimulation (Figure 4D) to awake mice while recording in the LCIC. Remarkably, ∼75% of LCIC neurons were activated by both high frequency mechanical vibration of the platform and auditory white noise, with a minority of neurons responding to only platform vibration or sound (Figure 4E-H). These findings reveal a high degree of convergence of auditory and mechanical vibration signals in the LCIC. Interestingly, of the LCIC neurons that received convergent tactile-auditory input, the majority exhibited a larger response to coincident presentation of mechanical vibration and sound than to either vibration or sound alone (Figure 4I-J).

We further assessed the multimodal nature of LCIC responses to environmental vibrations by measuring responses of LCIC neurons in mice lacking either Pacinian corpuscles or cochlear function (Figure 5A-B). As predicted from the findings in anesthetized mice (Figure 3D-G), LCIC neurons in awake Ret cKO mice, which lack Pacinian corpuscles, were unresponsive to high frequency surface vibrations (Figure 5E, G-H). Despite their insensitivity to vibration, LCIC neurons in Ret cKO mice were responsive to brush, suggesting that the LCIC receives inputs from other classes of cutaneous LTMRs. It is noteworthy that the somatosensory neuron deletion of Ret in Ret cKO mice does not impact the auditory system, as the Auditory Brainstem Response (ABR) test in Ret cKO mice indicated similar auditory thresholds to littermate controls (Figure S6A-C). Furthermore, LCIC neurons of Ret cKO mice were comparably sensitive to auditory white noise stimuli compared to controls, indicating that developmental loss of Pacinian corpuscles does not alter LCIC sensitivity to broadband auditory stimuli (Figure 5E, G-H).

**Figure 5.**
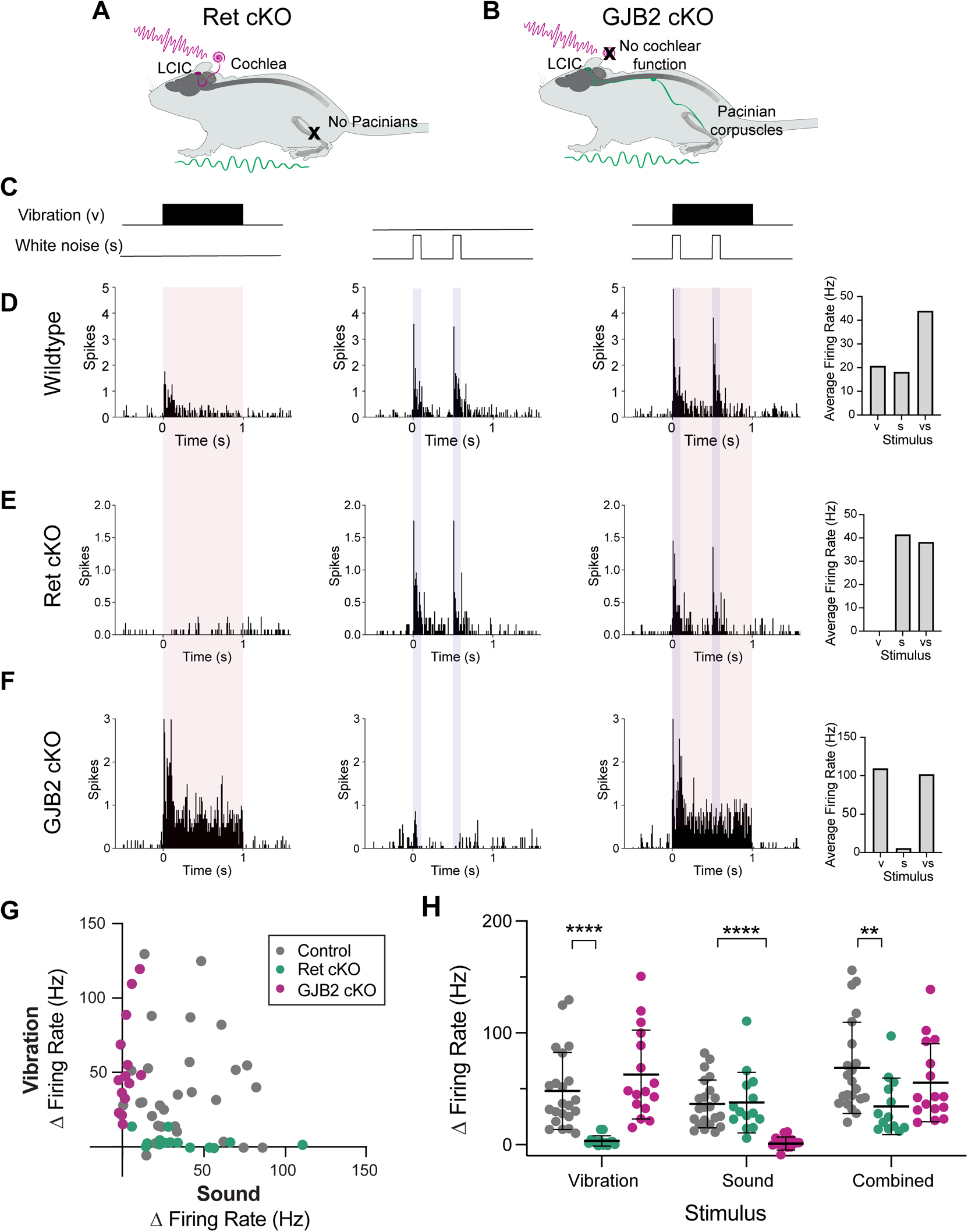
LCIC neuron responses in animals that lack Pacinian corpuscles or cochlear function. **A)** Schematic of Ret cKO mouse which lacks Pacinian corpuscles, but has intact hearing. **B)** Schematic of GJB2 cKO mouse which lacks cochlear function, but has an intact somatosensory system. **C)** 500 Hz substrate vibration (amplitude 0.2 g) and auditory white noise (frequency 0-50 kHz, 100 ms pulses, 65 dB SPL) were delivered in isolation and coincidentally to awake head-fixed mice. **D)** Example PSTHs from an LCIC neuron of a wildtype mouse aligned to vibration (v), sound (s) and convergent (vs) stimuli. PSTHs are baseline subtracted and in 10 ms bins. Right: The example neuron’s average firing (during 0-0.1 s and 0.5-0.6 s windows) for each stimulus type. **E and F)** As in (D), example LCIC neurons in a Ret cKO animal, which lacks Pacinian corpuscles (E), and a GJB2 cKO animal, which is unable to hear (F). **G)** Scatter plot of each neuron’s average change in firing rate to vibration and sound stimuli in control (n=29 neurons from 5 animals), Ret cKO (n=14 neurons from 3 animals) and GJB2 cKO (n=15 neurons from 3 mice) animals. **H)** Average change in firing rate to vibration, sound, or combined stimuli (mean ± SD) in control (n=29 neurons from 5 animals), Ret cKO (n=14 neurons from 3 animals) and GJB2 cKO (n=15 neurons from 3 mice) animals (**p<.01, ***p<0.001, ****p<0.0001, Mann-Whitney U test).

In complementary experiments, we tested whether disrupting auditory system function alters mechanical vibration responsiveness in the LCIC. For this, we utilized *Sox10^Cre^; GJB2^flox/flox^* mice (GJB2 cKO), which lack connexin 26 expression in epithelial cells of the cochlea^61,62^. GJB2 cKO mice lack cochlear hair cell function but have an intact vestibular system, such that this congenital deafness model exhibits extensive hearing loss but retains normal balance and motor function^61–65^. ABR testing in GJB2 cKO mice confirmed their inability to hear, as none of the GJB2 cKO mice tested had a discernable brainstem response at the highest sound level tested (Figure S6D-F). Recordings in anesthetized GJB2 cKO mice revealed that in the absence of normal hearing, LCIC neurons exhibit robust responses to mechanical vibration stimulation of the leg (Figure S6G-J). Force-frequency vibration tuning curves in GJB2 cKO showed no difference from controls (Figure S6I-J). Furthermore, recordings in awake GJB2 cKO mice demonstrated that LCIC neurons responded to substrate vibrations although they showed no responsivity to white noise auditory stimuli (Figure 5F-H). Thus, genetically disrupting either somatosensory/Pacinian corpuscle or auditory system function does not grossly alter LCIC neuron responses to the other sensory modality. Together, these findings indicate that most LCIC neurons receive both auditory input emanating from the cochlea and somatosensory input emanating from the body’s Pacinian corpuscles. Therefore, the LCIC is a site of auditory-mechanical vibration multisensory integration and enhancement.

### The LCIC mediates behavioral avoidance to tactile vibration

The LCIC has been implicated in mediating defensive behaviors in response to auditory cues^66^. To begin investigating whether the LCIC can drive behavioral responses to high frequency mechanical vibrations, we developed a behavioral preference assay that relies on an animal’s ability to detect and respond to high frequency mechanical vibrations of the environment. Animals were placed in a two-chamber behavioral setup where the floor of one chamber vibrates at 500 Hz, a stimulus that robustly activates Pacinian corpuscle afferents^15^ and most LCIC neurons of awake animals (Figure 4B-C, Figure 6A, Figure S7A). Accelerometers were attached to each chamber, confirming the consistency of the stimulus between trials as well as the containment of the vibration to one side of the chamber (Figure S7B-D). Wildtype animals had a strong and reliable preference for the non-vibrating chamber, spending ∼80% of their time there when the vibration stimulus was on (Figure 6B-D, Figure S7E-G, Supplementary Video 1). This avoidance was present in the two mouse strains tested, C57/Bl6 and CD-1 mice, and persisted in animals that were re-tested one week after the initial behavioral assay (Figure S7E-G). Furthermore, whisker trimmed animals showed a preference that was comparable to control animals, indicating that whiskers are not necessary for the behavior (Figure S7E-G). Since the production of the 500 Hz vibration generated a low amplitude 500 Hz auditory tone, we also performed a sound-only control. We decoupled the vibration motor from the chamber floor, such that a comparable sound was produced with no surface vibration. In the sound-only paradigm, animals showed no aversion toward the stimulus side of the chamber, demonstrating that sound alone does not drive a preference (Figure 6G-H). To test whether the high frequency mechanical vibration avoidance behavior is dependent on Pacinian corpuscles, the behavioral preference assay was done using Ret cKO mice, which lack Pacinian corpuscles. Ret cKO mice did not prefer the non-vibrating side of the chamber over the vibrating side, indicating that avoidance of the 500 Hz vibration stimulus is dependent on Pacinian corpuscles (Figure 6E-F, I-J).

**Figure 6.**
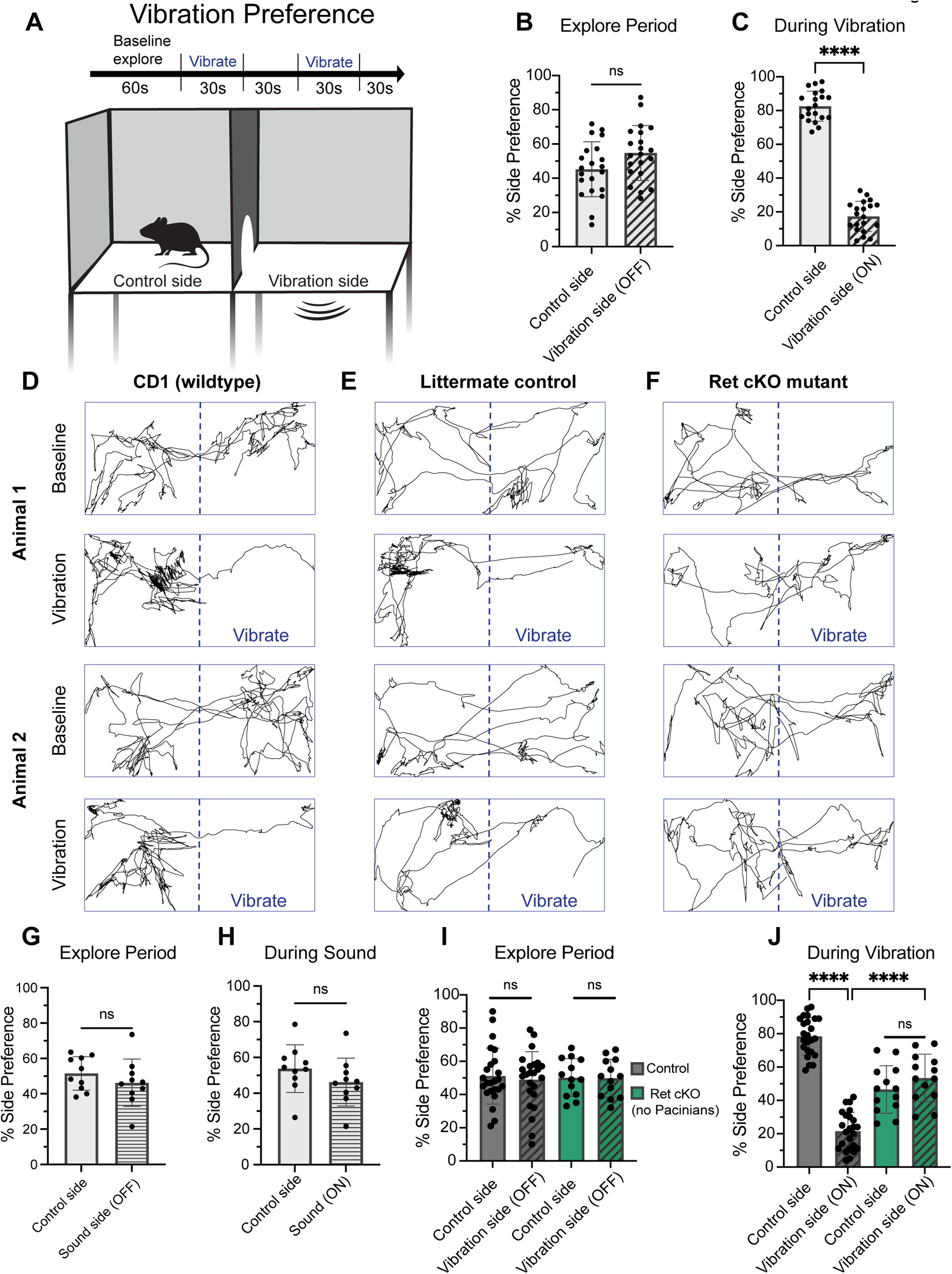
A behavioral paradigm to assess behavioral responsiveness to high frequency environmental vibration. **A)** Animals were placed in a 2-chamber behavioral setup. The floors of each chamber are disconnected to prevent propagation of vibration between them, and an exciter speaker is attached to the floor of one chamber to generate mechanical vibrations. The chamber walls are partially open to prevent sound resonance from the exciter. **B and C)** The percentage of time wildtype CD-1 animals spend on each side of the chamber (mean ± SD) during the 60 s baseline explore period (B) and the vibration periods (C) of the trial. Each dot represents one animal (n=20 animals, ****p<0.0001 paired t-test). **D-F)** Representative activity traces of two example wildtype CD-1 animals (D), littermate control animals (E), and Ret cKO animals (F), tracking movement during the baseline and vibration periods. **G and H)** The speaker was detached from the floor of the chamber to test whether the sound of the exciter speaker alone can drive a behavioral preference. Time spent on each side of the chamber during the 60 s baseline explore period (G) and the sound on periods (H) (not statistically different, n=10 animals, paired t-test). **I and J)** The percentage of time Ret cKO mice (n=14 animals) and littermate controls (n=24 animals) spent on each side of the chamber during the 60 s baseline explore period (I) and the vibration periods (J) of the behavioral trial. (****p<0.0001, paired t-test within group, unpaired t-test between groups).

The Pacinian corpuscle-dependent vibration place preference assay was next used to determine whether the behavioral response to 500 Hz mechanical vibration is dependent on the LCIC. Targeted, bilateral injections of a fluorescently tagged GABA-A receptor agonist, muscimol, were used to silence neural activity in the inferior colliculus (IC; Figure 7A, Figure S7H-J). Post-hoc histology confirmed that muscimol was reliably restricted to the IC and did not extend into neighboring brain regions, including the superior colliculus, periaqueductal gray, or cerebellum (Figure 7A, Figure S7H-I)^66–68^. IC-silenced mice displayed normal locomotor activity, traveling comparable distances during the assay as controls (Figure 7B). Strikingly, as with mice lacking Pacinian corpuscles, IC-silenced mice did not exhibit avoidance of the vibratory stimulus compared to saline injected controls, revealing that the IC is required for the behavioral response to high frequency mechanical vibration (Figure 7C-E, Supplementary Video 2). To determine whether the IC mediates general behavioral avoidance, or whether the effect of IC silencing is specific to vibratory stimuli, we also tested whether IC silencing alters avoidance of other somatosensory stimuli. We first performed a texture preference assay in saline and muscimol treated mice. In a two-chamber setup, one side of the chamber was covered in a smooth construction paper, while the other side was covered with a coarse grit sandpaper, which is aversive (Figure 7F, Figure S7K-L). Mice injected with either saline or muscimol in the IC showed strong preference for the smooth side of the chamber, revealing that the IC is not necessary for aversive texture avoidance (Figure 7G). A cold temperature avoidance test was also performed. For this, we used a two-chamber cold avoidance assay in which mice were given a choice between one chamber, which was held at room temperature (30 °C), and a second chamber, which was held at 18 °C (Figure 7H, Figure S7M). Similar to the texture preference assay, both control and IC-silenced animals displayed robust avoidance of the cold chamber (Figure 7I). These findings indicate that the IC mediates behavioral response to high frequency mechanical vibrations but is not required for avoidance of an aversive texture or temperature.

**Figure 7.**
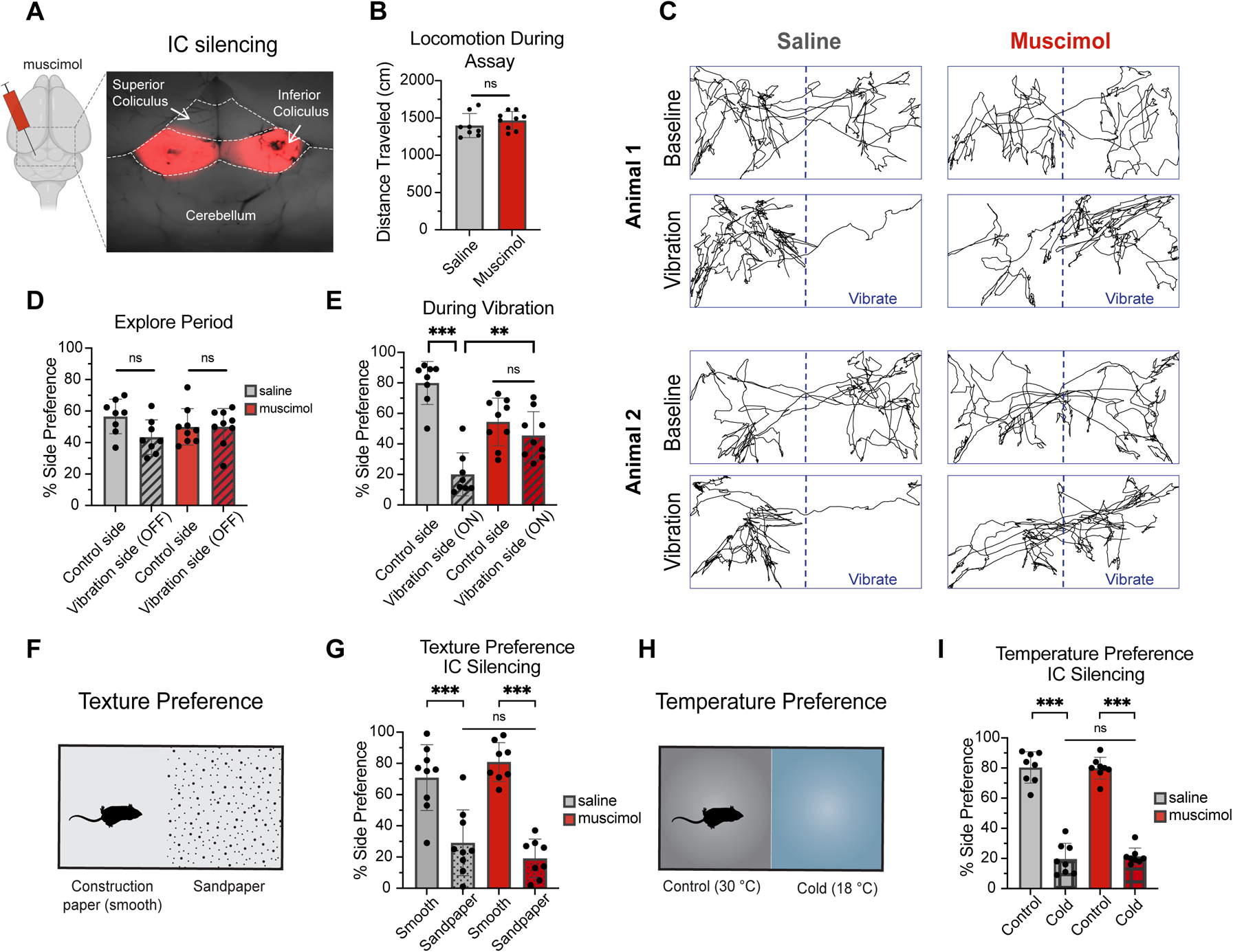
The LCIC mediates behavioral avoidance of high frequency mechanical vibration. **A)** Targeted injections of TMR-X conjugated to the GABA-A receptor agonist, muscimol, were used to silence neural activity in the inferior colliculus. Top-down view of a schematic (left) and representative muscimol injected brain. Dotted lines depict borders between the inferior colliculus and indicated neighboring brain regions. **B)** Total distance traveled during the behavioral assay in saline injected controls (n=8) and muscimol injected animals (n=9 animals, not significant, unpaired t-test). **C)** Representative activity traces for 2 saline injected (control) and 2 muscimol injected animals during the baseline and vibration periods of the behavioral trial. **D and E)** The proportion of time muscimol injected mice (n=9 animals) and saline injected controls (n=8 animals) spent on each side of the chamber during the 60 s baseline explore period (D) and the vibration periods (E) of the trial. (**p<0.01, ***p<0.001, paired t-test within group, unpaired t-test between groups) **F)** Schematic of texture preference assay. **G)** Percentage of time spent on the smooth and rough sides of the chamber during the 10 min assay for saline controls (n=9 animals) and muscimol silenced animals (n=9 animals, ***p<0.001, paired t-test within group, unpaired t-test between groups). **H)** Schematic of temperature preference assay. **I)** Percentage of time spent on the control (30 °C) and cold (18°C) sides of the chamber during the 5 min assay for saline controls (n=8) and muscimol silenced animals (n=8, ***p<0.001, paired t-test within group, unpaired t-test between groups).

## Discussion

Organisms across the animal kingdom, including insects, reptiles and mammals, rely on their ability to detect and respond to environmental vibratory cues for their survival. Spider webs are designed to transmit vibrations that spiders utilize to efficiently localize their prey^5^. Snakes detect the movement of prey by pressing their jaws into the ground to detect subtle substrate vibrations^6^. Some subterranean mammals, in the absence of visual cues, rely heavily on vibratory cues for communication, localization of prey, and detection of predators^1–3^. While all mammalian LTMRs are sensitive to mechanical stimuli acting on the body, the Pacinian corpuscle is unique in its exquisite sensitivity to substrate vibrations of the environment and across a broad range of frequencies. Recordings from Pacinian afferents in awake, behaving mice demonstrate their capacity to fire robustly in response to subtle vibrations generated meters away from where an animal stands^15^. Here we show that the neural signals encoding mechanical vibrations captured by the body’s Pacinian corpuscles converge with the signals encoding auditory vibrations (sound) in the LCIC of the auditory midbrain to create an internal representation of environmental vibration.

### Divergent functions of thalamic and collicular pathways

The majority of research into the neurobiological basis of discriminative touch of the body has focused on the ascending pathway through the brainstem to the thalamus (VPL) *en route* to primary somatosensory cortex (S1; Figure 1A), while somatosensory representations in other brain areas have remained largely unexplored. We found that neurons in the VPL and the LCIC of the auditory midbrain exhibit highly distinct vibrotactile tuning properties that may underlie unique functions of divergent ascending somatosensory system streams. Most VPL neurons had restrictive receptive fields and were tuned to low frequency vibrations. These properties are well suited for tactile acuity and discrimination—functions hypothesized to be mediated by the VPL to S1 pathway^69–72^. Only a small fraction of VPL neurons (∼9%) encoded high frequency vibration (Figure 1D). In contrast, most LCIC neurons exhibited expansive receptive fields and were tuned to high frequency vibrations (>75%) conveyed by Pacinian corpuscle Aβ RA2-LTMRs (Figure 1D, F). These divergent somatosensory streams are reminiscent of the parallel processing streams of the mammalian visual system, where some retinal ganglion cells project to the thalamus (LGN), while others project to the tectum (superior colliculus) or the suprachiasmatic nucleus (SCN) of the hypothalamus^73^. Work in the visual system suggests that functional segregation exists between the LGN, tectal, and SCN pathways, whereby the collicular pathway may be optimally tuned for discrimination of moving objects^74^ and the SCN pathway for circadian entrainment^73^. These parallels between visual and somatosensory processing reveal shared organizational principles across sensory systems.

The salience of low amplitude environmental vibrations encoded in the LCIC is highlighted by our finding that IC silencing virtually eliminated animals’ avoidance of a 500 Hz vibratory stimulus in a place preference paradigm (Figure 7C-E). On the other hand, the IC was not required for avoidance of an aversive texture or temperature, suggesting that the IC does not mediate general avoidance to aversive somatosensory stimuli, rather it plays a more specialized role in perceiving and reacting to high frequency vibrations (Figure 7G,I). It is possible that the divergent ascending streams through the LCIC and VPL underlie distinct responses to high frequency vibrations in different contexts. For example, the sparser representation of high frequency vibration in the VPL to S1 stream may underlie operant tasks that use vibratory cues^14,71,72,75^. Future work will discern the relative contributions of distinct ascending streams encoding environmental vibrations in different contexts and downstream targets of the LCIC that underlie environmental vibration sensing.

### Tactile-auditory integration in the LCIC

Strikingly, most mechanical vibration sensitive LCIC neurons are also tuned to auditory stimuli. Moreover, the majority of LCIC neurons that received convergent mechanical vibration and auditory inputs exhibited multisensory enhancement, responding more strongly to coincident tactile-auditory stimulation than to either modality alone (Figure 4I-J). Furthermore, somatosensory and auditory inputs to the LCIC are independent of one another, as evidenced by the findings that genetically altering the ability to detect either mechanical or auditory inputs did not grossly impact responsivity to the other modality (Figure 5). Thus, the LCIC encodes environmental vibrations, whether they are mechanical vibrations propagated through the ground, detected by Pacinian corpuscles, and conveyed via the ascending somatosensory pathway or sound waves traveling through air, captured by the cochlea, and conveyed via the ascending auditory pathway.

We suspect that the LCIC enables enhanced detection or reaction to environmental vibrations through auditory and vibrotactile coincidence detection. A myriad of natural phenomena generates coincident sounds and substrate vibrations, including seismic events or earth movements, objects moving across or falling to the ground, vocalizations of conspecifics, and the motions of prey or predators. Multisensory audio-vibrotactile integration in the LCIC may increase saliency of these ethologically relevant environmental cues and may be particularly relevant under conditions in which cues are weak and difficult to discern. For humans, the connection between vibration and sound is evident when standing in the wind, close to a rushing river or a passing train, in a concert, or playing a musical instrument. We propose that, through their convergent inputs to the LCIC, the somatosensory and auditory systems collaborate to detect, localize, and respond to a wide range of environmental vibrations.

**Supplementary Figure 1.**
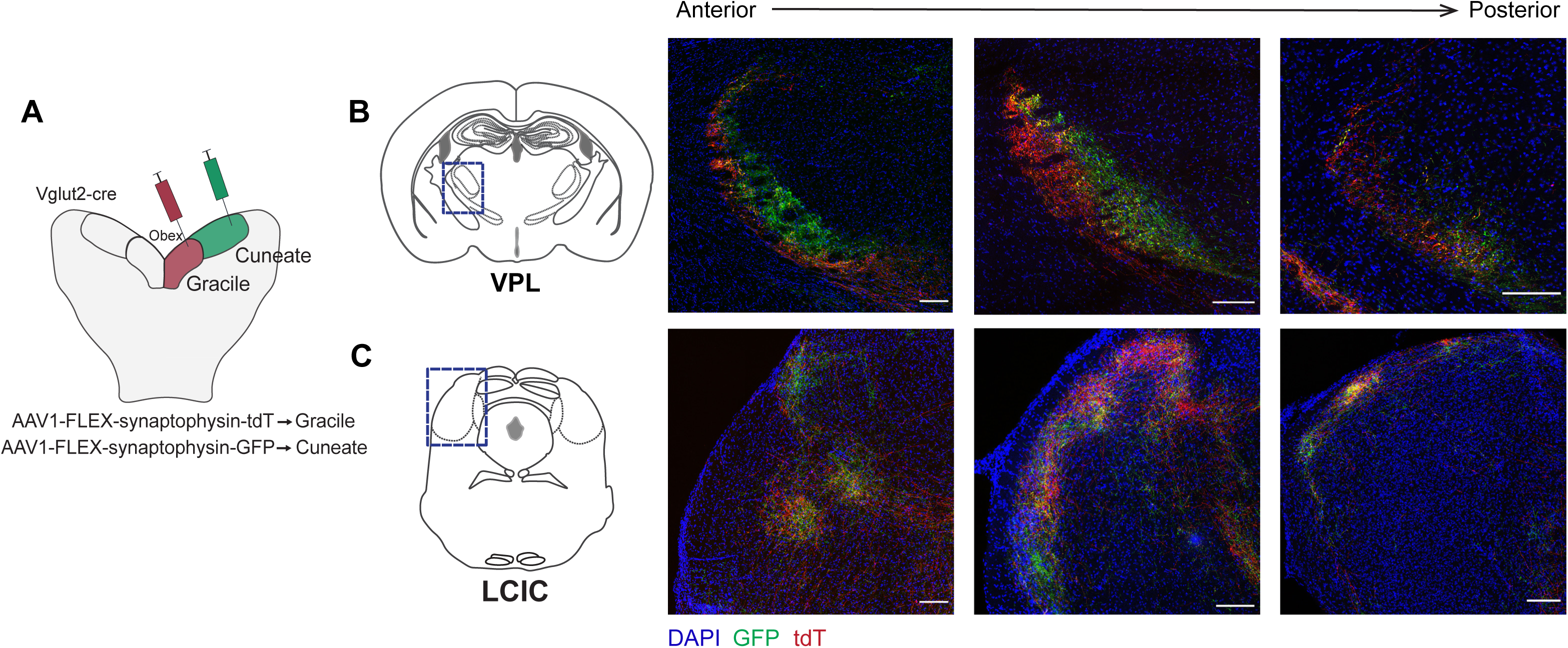
Anterograde tracing from the dorsal column nuclei. **A)** To identify downstream projections from the dorsal column nuclei (DCN), an AAV encoding Cre-dependent synaptophysin-tdT was injected into the gracile nucleus and an AAV encoding Cre-dependent synaptophysin-GFP was injected into the cuneate nucleus of Vglut2-Cre mice. **B-C)** TdT and GFP positive terminals were observed throughout the ventroposterolateral thalamus (VPL) and lateral cortex of the inferior colliculus (LCIC). Scale bar = 150 μm.

**Supplementary Figure 2.**
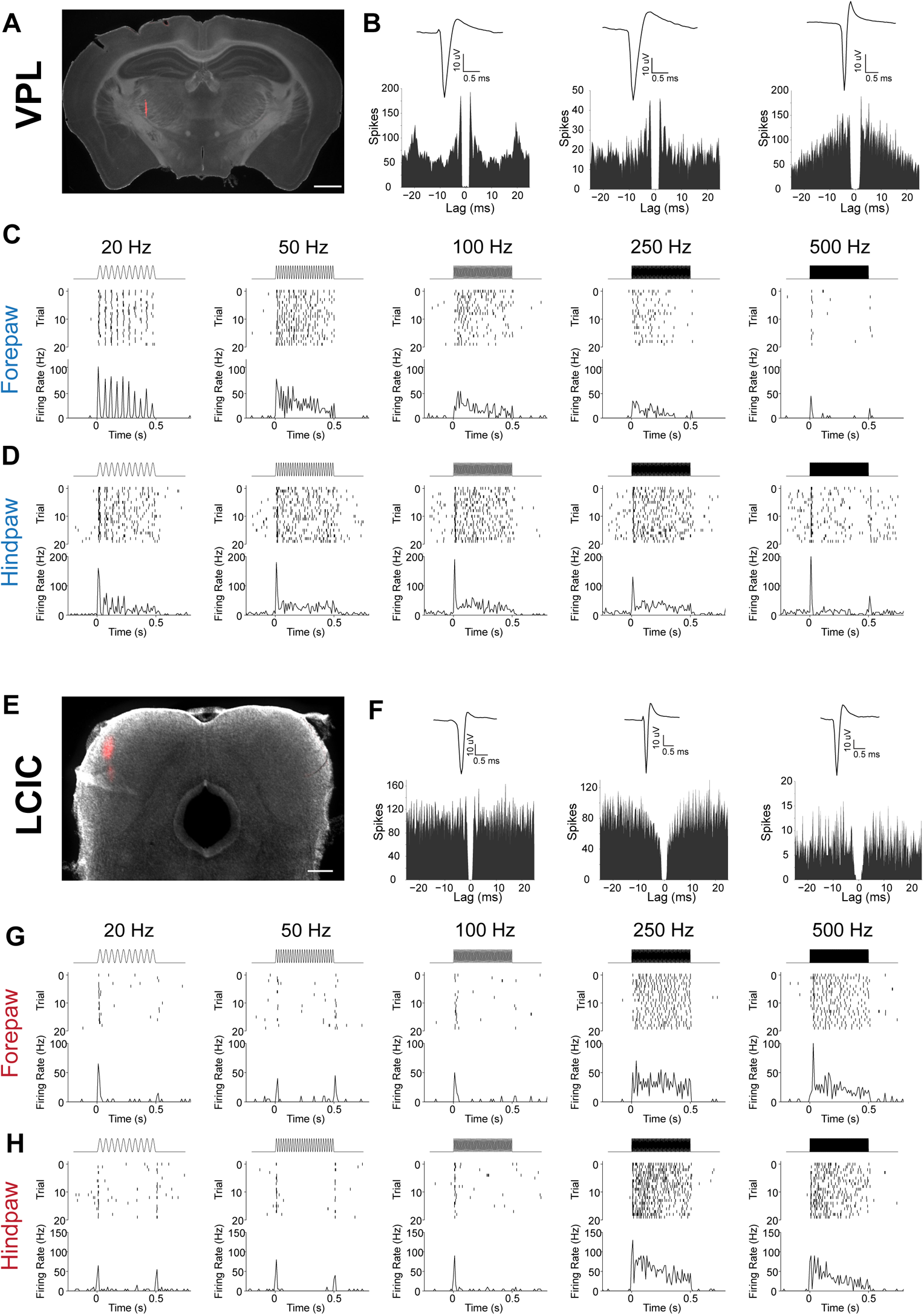
MEA recordings in the VPL and LCIC to characterize vibratory responses. **A)** Representative probe tract from MEA recording in the VPL. Scale bar = 1000 μm. **B)** Average waveforms and corresponding autocorrelogram for three example VPL neurons. **C)** Example VPL neuron raster and PSTHs in response to vibration of different frequencies delivered at a 10 mN force to the forelimb. Top: spike raster; bottom: PSTHs, which are baseline subtracted and in 10 ms bins. **D)** Same as C for a hindlimb responsive neuron. **E)** Representative probe tract from MEA recording in the LCIC. Scale bar = 500 μm. **F)** Average waveforms and corresponding autocorrelogram for three example LCIC neurons. **G)** Same as C for an example LCIC neuron. **H)** Same as D for an example LCIC neuron.

**Supplementary Figure 3.**
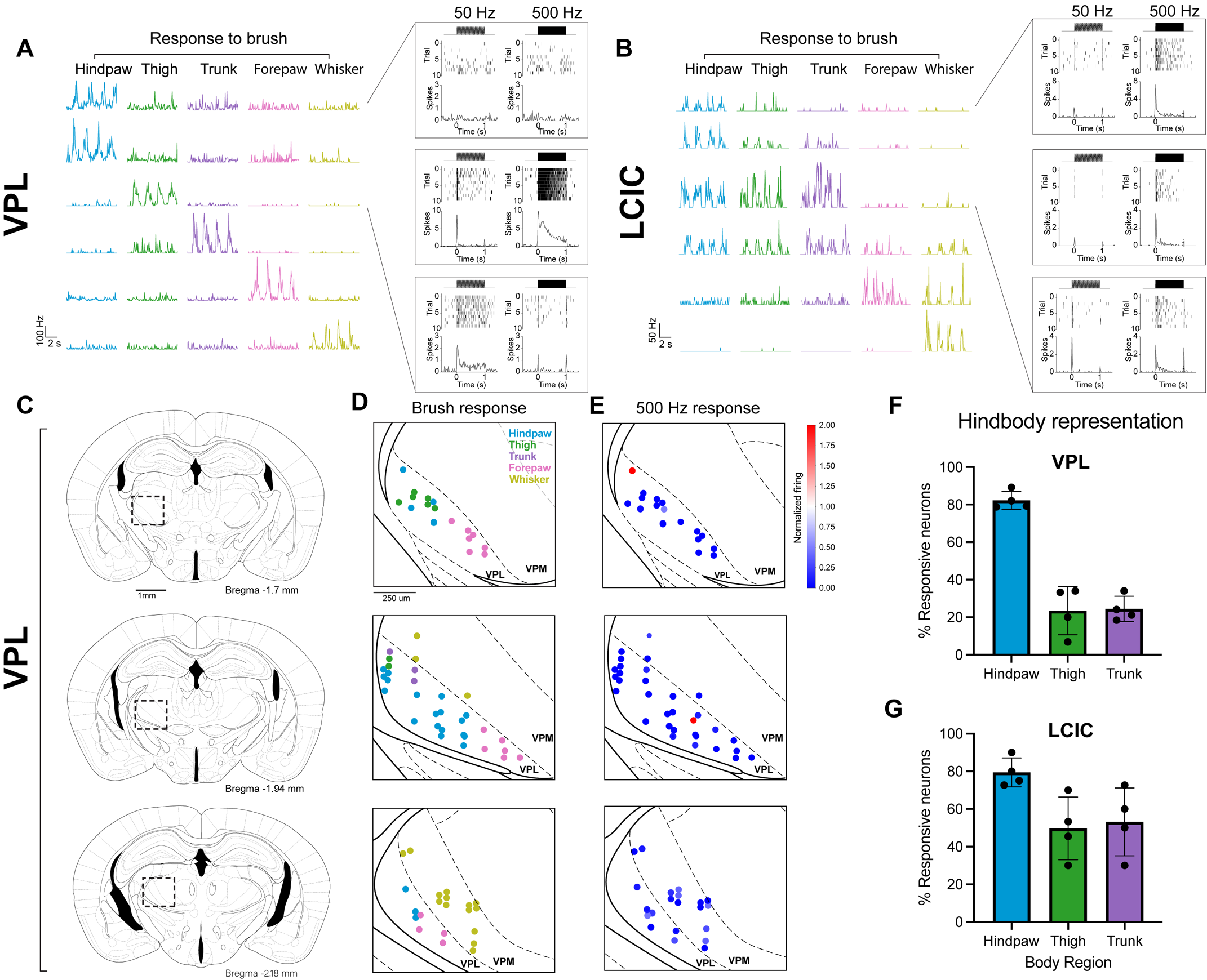
Somatotopic and vibratory mapping in the VPL and LCIC. **A)** Representative VPL neurons responding to brush of different regions of the body. Each row corresponds to one neuron. Each body region is stroked four times per trial. Right: Example raster plot and PSTHs aligned to hindlimb vibration at 50 Hz and 500 Hz at 10 mN, corresponding to adjacent brush responsive neurons. PSTHs are baseline subtracted and in 10ms bins. **B)** As in A, representative LCIC neurons. Note that, unlike VPL neurons, most LCIC neurons respond to brush across multiple body locations. **C-E)** Example mapping of the VPL in one animal, shown at different coronal planes on the anterior-posterior axis. The approximate atlas location (C) for recorded neurons is shown for the corresponding somatotopic map of the VPL (D), where each neuron is colored for the body region most responsive to brush. The same neurons shown in D are colored by their responsiveness to 500 Hz mechanical vibration, which is normalized to the neuron’s response to brush (E). Note that most brush responsive VPL neurons do not respond to the 500 Hz vibration stimulus during the sustained period of the stimulation. **F and G)** Percentage of neurons (mean ± SD) with hindbody receptive fields that respond to brush of the hindpaw, thigh, and trunk in the VPL (F) (n= 182 neurons from 4 animals) and LCIC (G) (n= 76 neurons from 4 animals). Each data point corresponds to a single animal.

**Supplementary Figure 4.**
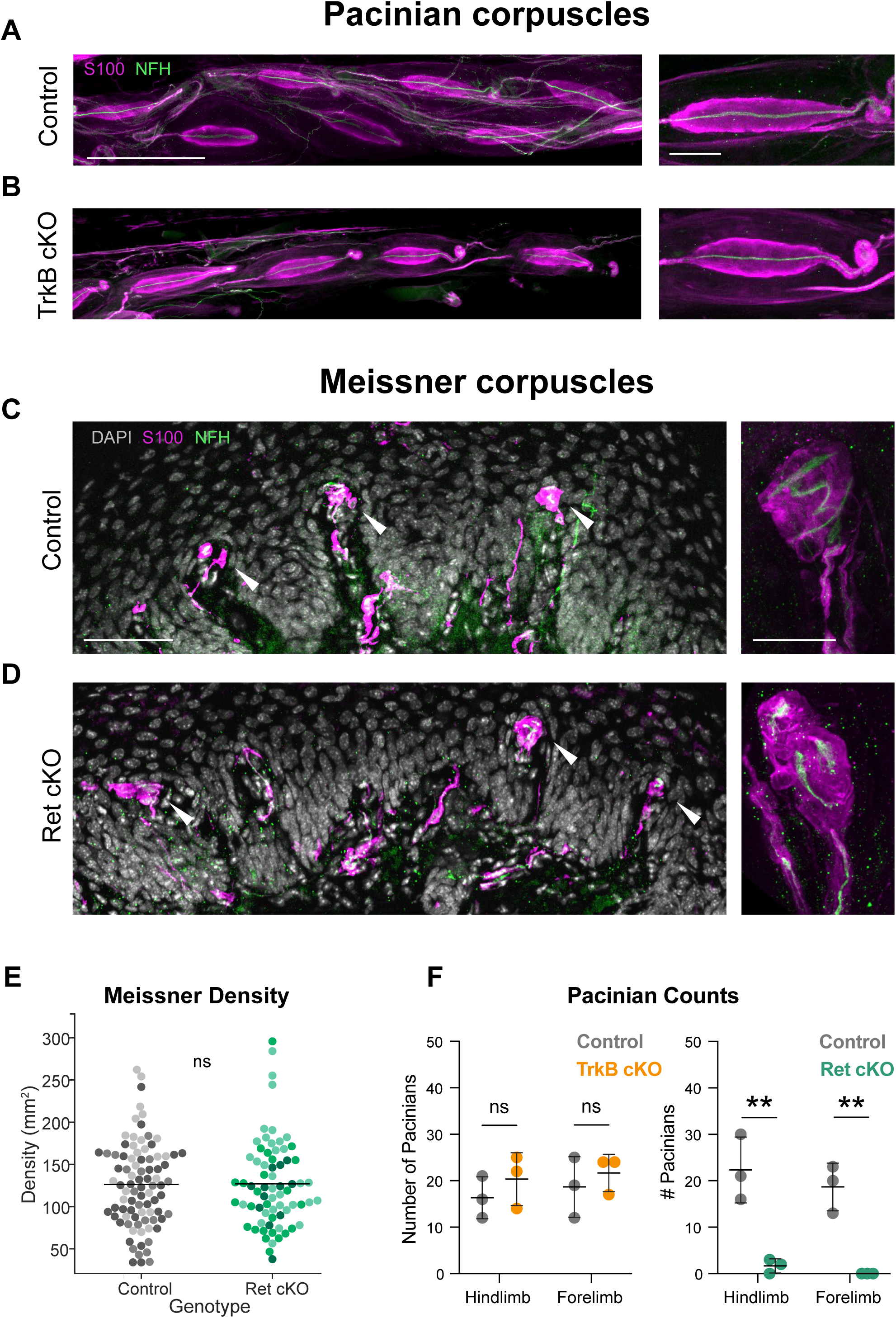
Anatomical characterization of LTMR peripheral endings in Ret cKO and TrkB cKO mice. **A-B)** Images of Pacinian corpuscles from the limbs of TrkB cKO animals and littermate controls. Limbs were whole-mount stained with antibodies against NFH (green), which labels large caliber axons, and S100 (magenta), which labels lamellar cells of the corpuscle and myelinating Schwann cells. Left scale bar = 250 μm. Right scale bar = 50 μm. **C-D)** Images of Meissner corpuscles (indicated by white arrows) from Ret cKO animals and littermate controls. Skin tissue was stained using antibodies against NFH (green) and S100 (magenta). DAPI staining is shown in gray. Left scale bar = 50 μm. Right scale bar =20 μm. **E)** Quantification of Meissner density in Ret cKO (n=3 animals) and littermate controls (n=3 animals). Each data point is the Meissner density for a single skin section, and data points of the same color came from the same animal (no significant difference using unpaired t-test). **F)** Quantification of Pacinian corpuscles in TrkB cKO (n=3 animals) and littermate controls (n=3 animals), and Ret cKO (n=3 animals) and littermate controls (n=3 animals). The number of Pacinians was counted from wholemount staining of hindlimbs and forelimbs (mean ± SD, **p<0.001, unpaired t-test).

**Supplementary Figure 5.**
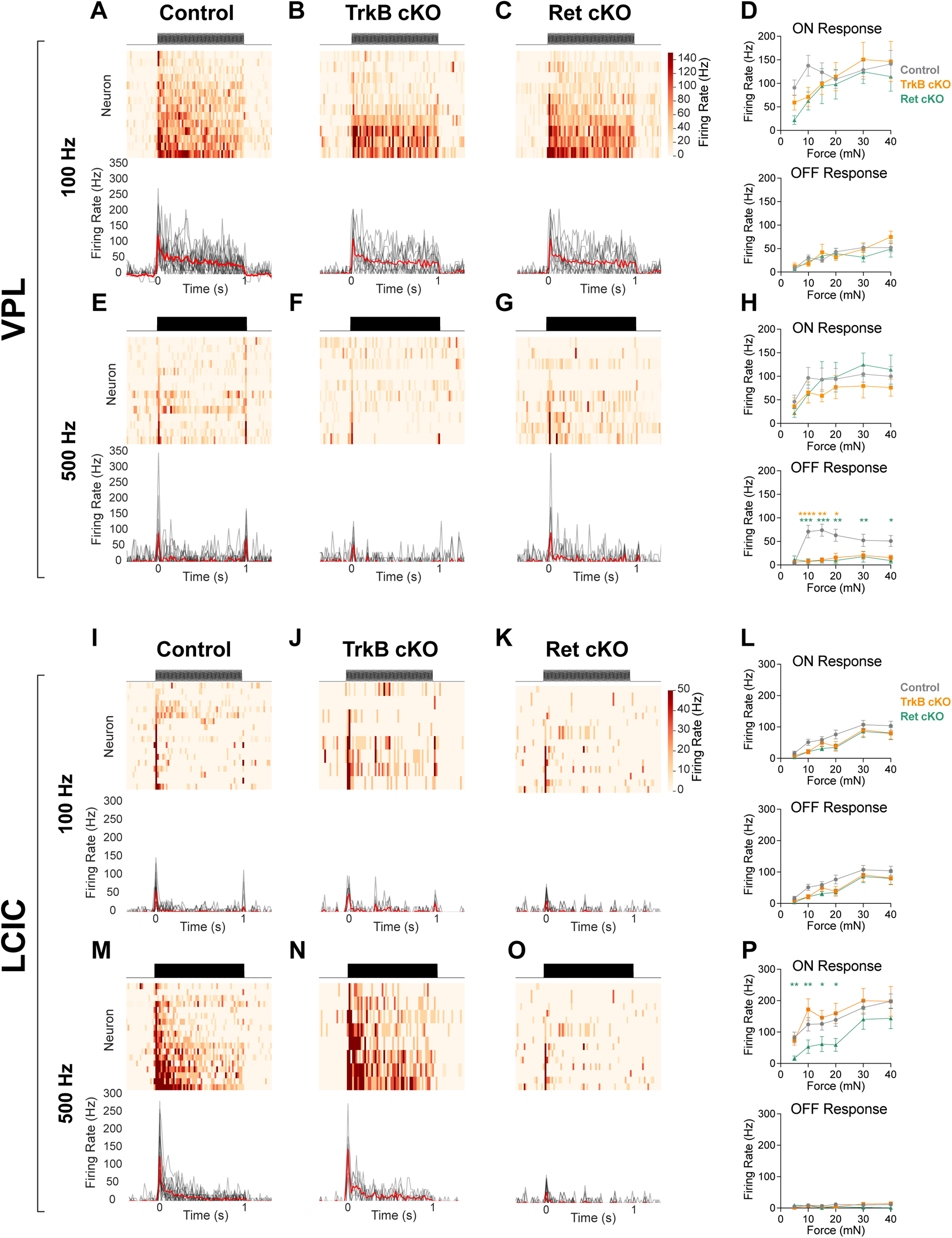
Onset and offset responses to vibratory stimuli in Ret cKO and TrkB cKO mice. **A)** Top: Heatmap of neurons in the VPL responding to 100 Hz vibration at a 15 mN force (n=14 neurons from 6 animals). Each row is one neuron. Bottom: PSTHs of each neuron in above heatmap (gray), and population average PSTH (red). PSTHs are in 20ms bins. **B-C)** Same as A in TrkB cKO (n=10 neurons from 3 animals) and Ret cKO mice (n=14 neurons from 4 animals). **D)** Average firing rate of VPL neurons to the onset (top) and offset (bottom) of the 100 Hz vibration stimulus in control, TrkB cKO, and Ret cKO mice (mean ± SEM, not significantly different, Mann-Whitney U test with Bonferroni correction for multiple comparisons). On period is 20 ms after the stimulus onset, and off period is 20 ms after the end of the stimulus. **E-G)** Same as A-C to 500 Hz vibration at 15 mN. **H)** Same as D for 500 Hz stimulus (*p<0.05, **p<0.01, ***p<0.001, ****p<0.0001, Mann-Whitney U test with Bonferroni correction for multiple comparisons). **I-K)** Same as A-C for neurons in the LCIC in control (n=18 neurons from 5 animals), TrkB cKO (n=8 neurons from 4 animals), and Ret cKO (n=16 neurons from 5 animals) animals. **L)** Average firing rate of LCIC neurons to the onset (top) and offset (bottom) of the 100 Hz vibration stimulus in control, TrkB cKO, and Ret cKO mice (mean ± SEM, not significantly different, Mann-Whitney U test with Bonferroni correction for multiple comparisons). **M-O**) Same as I-K to 500 Hz vibration at 15 mN. **P)** Same as L for a 500 Hz stimulus (*p<0.05, **p<0.01, Mann-Whitney U test with Bonferroni correction for multiple comparisons).

**Supplementary Figure 6.**
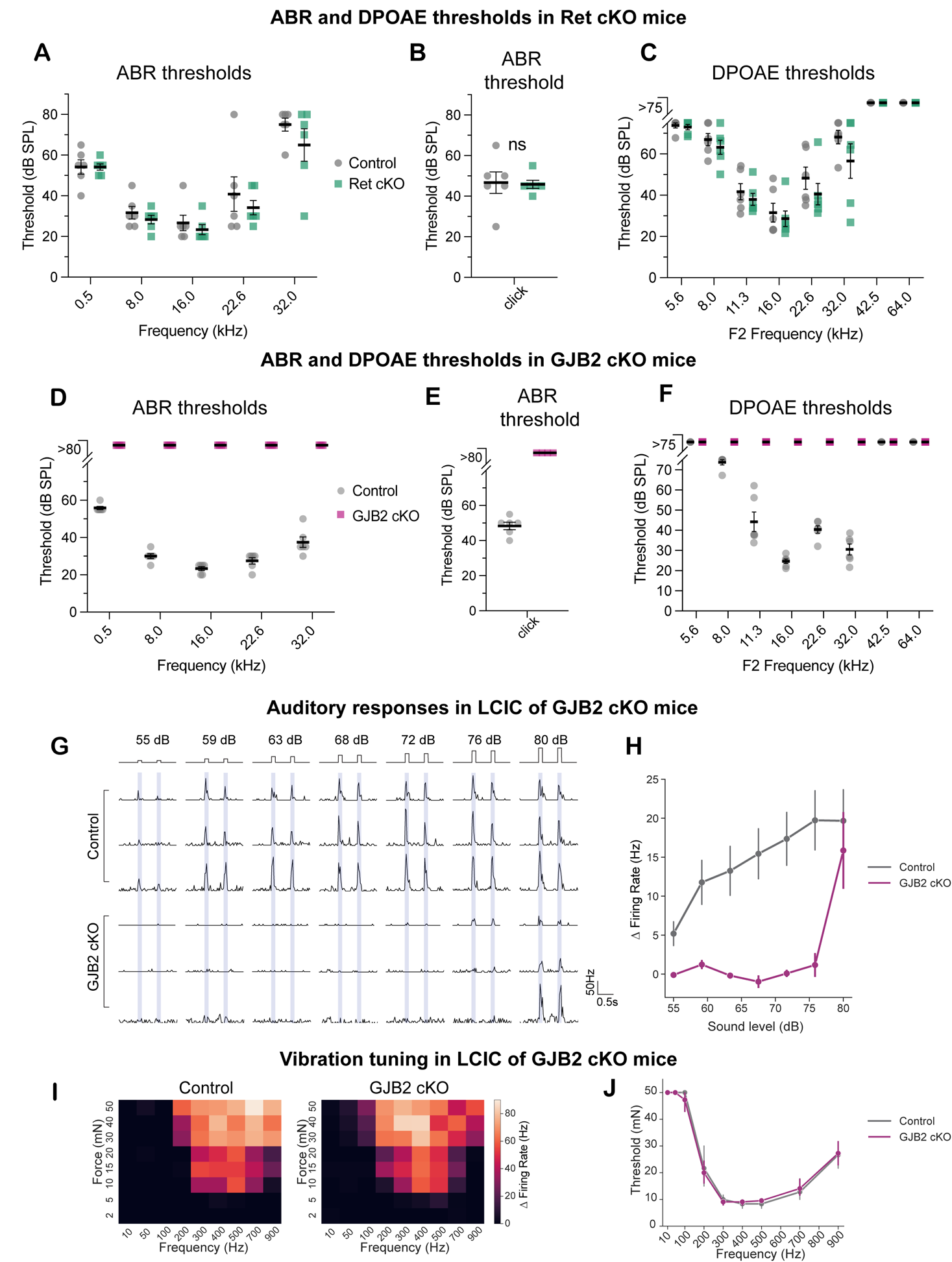
Characterization of auditory and tactile responses in Ret cKO and GJB2 cKO mice. **A)** Auditory brainstem response (ABR) thresholds were measured in Ret cKO mice and littermate controls at 5 different auditory frequencies (mean ± SEM). Auditory thresholds between Ret cKO (n=6 animals) and controls (n=6 animals) were not significantly different (Mann-Whitney U Test with Bonferroni correction for multiple comparisons). **B)** ABR thresholds to a broadband frequency “click” were measured in Ret cKO mice (n=6 animals) and littermate controls (n=6 animals). Thresholds were not significantly different (Mann-Whitney U Test). **C)** Distortion product otoacoustic emissions (DPOAE) thresholds, which are sounds generated by mechanically active cells in the ear in response to sound stimuli, were measured in Ret cKO mice (n=6 animals) and littermate controls (n=6 animals) for 8 different frequency pairs (mean ± SEM). Thresholds were not significantly different (Mann-Whitney U Test with Bonferroni correction for multiple comparisons), suggesting normal function of the sensory cells responsible for fine-tuning cochlear gain. **D)** Auditory brainstem response (ABR) thresholds were measured in GJB2 cKO mice (n=4 animals) and littermate controls (n=6 animals) at 5 different auditory frequencies (mean ± SEM). Unlike littermate controls, GJB2 cKO animals had no response at any frequency, even at the highest sound level tested (80 dB). **E)** ABR thresholds to a broadband frequency “click” were measured in GJB2 cKO mice (n=4 animals) and littermate controls (n=6 animals). GJB2 cKO animals had no response, even at the highest sound level tested (80 dB). **F)** Distortion product otoacoustic emissions (DPOAE) thresholds were measured in GJB2 cKO mice (n=4 animals) and littermate controls (n=6 animals) for 8 different frequency pairs (mean ± SEM). GJB2 cKO animals had no DPOAE at any frequency, even at the highest sound level tested (75 dB). **G)** Example neurons responding to white noise at varying sound levels (dB) in anesthetized littermate control and GJB2 cKO mice. Each row corresponds to an individual neuron. **H)** Average change in firing rate of neurons (mean ± SE) in littermate control (n=13 neurons from 2 mice) and GJB2 cKO mice (n=9 neurons from 2 mice) to white noise at varying sound intensities. **I)** Force-frequency heat maps in response to vibration of differing frequencies and intensities in a representative neuron from a littermate control and GJB2 cKO animal. **J)** Average force-frequency threshold curves of LCIC neurons (mean ± 95% confidence interval) recorded in control (n=9 neurons from 2 animals), and GJB2 cKO mice (n=11 neurons from 2 animals). Thresholds between GJB2 cKO and controls were not significantly different (Mann-Whitney U Test with Bonferroni correction for multiple comparisons).

**Supplementary Figure 7.**
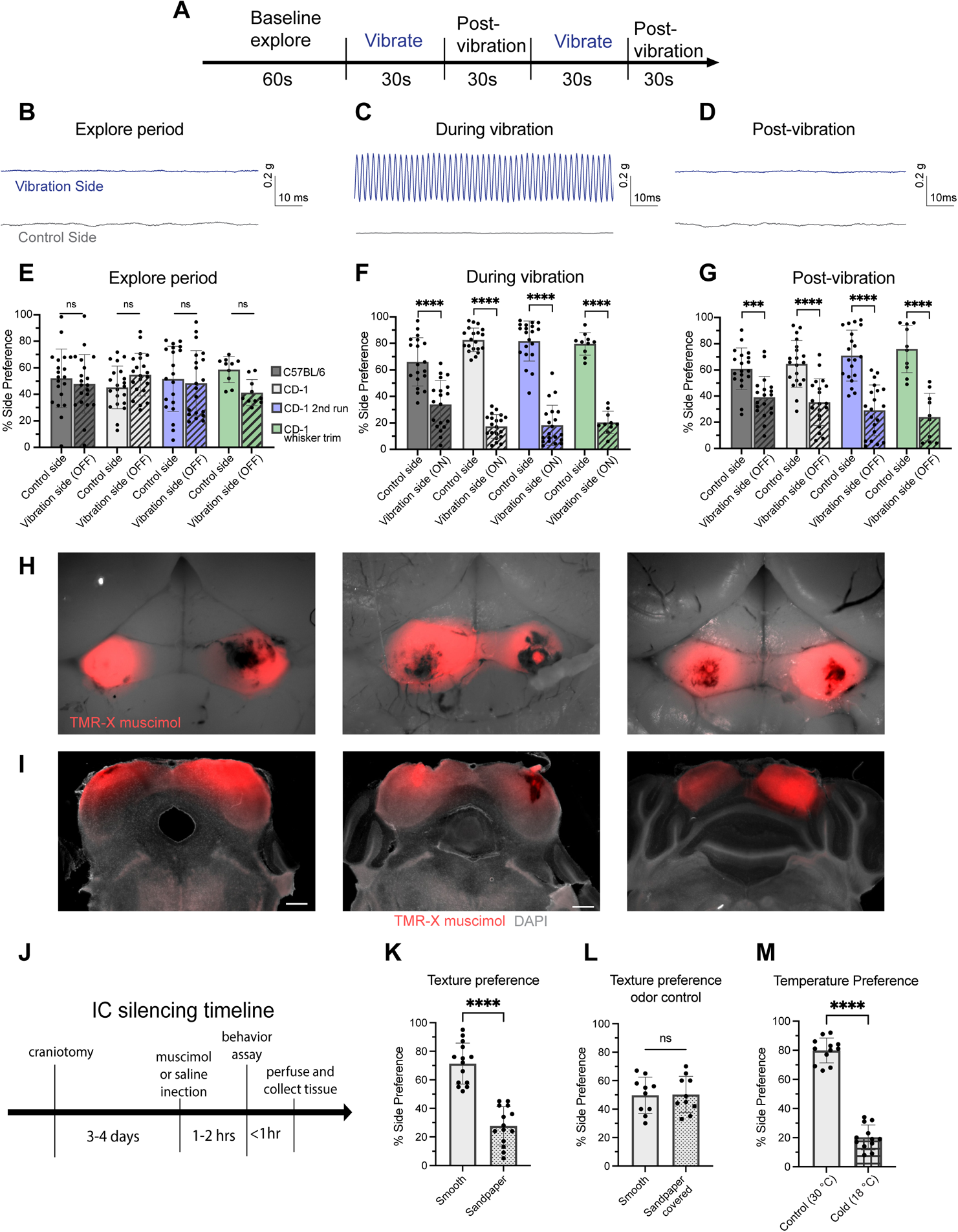
Behavioral analyses in wildtype and LCIC silenced animals. **A)** Timeline of behavioral trial. **B-D)** Accelerometer traces from vibration side (blue) and control side (gray) of the behavioral chamber during the baseline period (B), vibration period (C), and post-vibration period (D) of the behavioral trial. Amplitude of the platform vibration was ∼ 0.3 g (gravitational force) when the vibration stimulus was on. **E-G**) Percent of time that mice spent on the control and vibration side of the chamber during the explore period (E), vibration period (F), and post-vibration period (G) of the behavioral trial for four different conditions: naïve C57BL/6 mice (n=19 animals), naïve CD-1 mice (n=20 animals), CD-1 mice run on the behavior for the second time, one week later (n=20 animals), and naïve CD-1 mice that had been whisker trimmed (n=10 animals; ***p<0.001, ****p<0.0001, paired t-test). **H)** Post-hoc histology of three different TMR-X conjugated muscimol injected animals, showing restriction of the injection to the inferior colliculus. **I)** Coronal sections of a representative TMR-X conjugated muscimol injected brain to show ventral spread of the injection. Scale bar = 500 μm. **J)** Timeline of IC silencing experiments. **K)** Texture preference assay done for naïve CD-1 animals (n=10 animals, ****p<0.0001, unpaired t-test). Note, this is a separate cohort from the group used in silencing experiments in Figure 7. **L)** Some types of sandpaper are aversive to mice due to their odor. To ensure our texture avoidance behavioral measure reflects avoidance of an aversive texture and not odor, the sandpaper was covered by a smooth paper with small holes punched in it, such that the odor of the paper could be sensed, but the texture could not be felt. In this condition, mice (n=10 animals) did not demonstrate an aversion to the sandpaper side. (Not significant, paired t-test.) **M)** Temperature preference assay done for naïve wildtype CD-1 animals (n=10 animals, ****p<0.0001, unpaired t-test). Note, this is a separate cohort from the group used in silencing experiments, reported in Figure 7.

## Methods

### Animals

*Avil^Cre^*, *TrkB^flox^*, *Ret^flox(eGFP)^*, and *Vglut2^Cre^* mouse lines were of mixed background (CD-1 and C57BL/6J). *GJB2^flox^* and *Sox10^Cre^* were on a C57BL/6 background. CD-1 animals were imported from Charles River (#022) and C57BL/6 animals were imported from The Jackson Laboratory (#000664). The following mouse lines have been previously described, including: *Sox10^Cre^* (Ref 78, Jax 025807), *Avil^Cre^*(Ref: 79, JAX 032536), *TrkB^flox^* (Ref:80), *Ret^flox(eGFP)^* (JAX 029846), and *Vglut2^Cre^* (JAX 016963). The *GJB2^flox^* mouse line, in which loxP sites flank the entire coding sequence in exon 2, was generated by KTB in the laboratory of DPC and will be described elsewhere. Mice were handled and housed in standard cages in accordance with Harvard Medical School and IACUC guidelines. Both male and female mice were used in all experiments. Control littermates of *Ret*, *TrkB,* and *GJB2* conditional mutants included Cre-negative animals that were negative, heterozygous, or homozygous for the floxed allele. Our data indicated no difference among controls of different genotypes; therefore, they were combined for statistical analyses and collectively referred to as “controls.”

### Perfusions

Mice were anesthetized under isoflurane and transcardially perfused with 5-10 mL of PBS, followed by 5-10 mL of 4% paraformaldehyde (PFA) in PBS. After perfusion, the skull was removed and post-fixed in 4% PFA in PBS at 4°C overnight. Skin samples were removed and fixed overnight in Zamboni’s fixation buffer with gentle agitation at 4°C. Samples were washed 3x10 min in PBS at room temperature after overnight fixation before fine dissection.

### Immunohistochemistry of skin cryosections, brain cryosections and whole-mount tissue

The following dilutions and primary antibodies were used: chicken anti-NFH (Aves # NFH, 1:500), rabbit anti-S100 Beta (VWR/Protein Tech #15146-1-AP, 1:300), rabbit anti-dsRed (Takara #632496. 1:1000), chicken anti-GFP (Abcam #13970, 1:1000), goat anti-GFP (US Biological Life Sciences # G8965-01E, 1:500). The secondary antibodies used were Alexa 488, 546 or 647 conjugated donkey or goat anti-rabbit, chicken, or goat (Life Technologies or Jackson ImmunoResearch) and were prepared at 1:500 or 1:1000 dilutions.

For Meissner corpuscle histology, paws were removed from perfused mice. The pads and fingertips were clipped off of the glabrous skin using small spring scissors and cryoprotected in 30% <u>sucrose</u> in PBS at 4 °C for 1-2 days, embedded in OCT (1437365, Fisher), frozen using dry ice, and stored at −80 °C. The tissues were cryosectioned (25 μm) using a <u>cryostat</u> (Leica) and collected on glass slides (12-550-15, Fisher). Sections were washed 3 times for 5 minutes each with PBS containing 0.1% Triton X-100 (0.1% PBST), incubated with blocking solutions (0.1% PBST containing 5% normal goat serum (S-1000, Vector Labs) or normal donkey serum (005-000-121, Jackson Immuno) for 1 hour at room temperature, incubated with primary antibodies diluted in blocking solutions at 4 °C overnight, washed 3 times for 10 minutes each with 0.1% PBST, incubated with secondary antibodies diluted in blocking solutions at 4 °C overnight, washed again 4 times for 10 minutes each with 0.1% PBST (DAPI solution was included in the second wash at 1:5000 dilution), and mounted in Fluoromount-G mounting medium (Fisher 0100-01).

For whole-mount staining of Pacinian corpuscles, animals were anesthetized under isoflurane and euthanized by cervical dislocation. For Pacinian corpuscle dissections, the tibia and fibula of the ankle, and radius and ulna of the wrists were dissected immediately after euthanasia, keeping muscle and surrounding tissue intact, and fixed overnight in Zamboni’s fixative at 4 °C. Limbs were washed 3x10 minutes in PBS and then finely dissected to remove muscle on the bone, while leaving the interosseous membrane of the bone intact. Tissues were then washed every 30 minutes for 5-6 hours with 1% PBST (PBS + triton X-100). Primary antibodies were applied in blocking solution (5% normal goat or donkey serum, 75% PBST, 20% DMSO, 0.01% NaN3) and incubated at room temperature for 3-5 nights with gentle rocking. Tissues were rinsed 3x in PBST, then washed every 30 minutes for 5-6 hours in 1% PBST. Secondary antibodies were applied in blocking solution and incubated at room temperature for 3-5 nights, with gentle rocking. Tissues were rinsed 3x in 1% PBST, washed every 30 minutes for 5-6 hours in 1% PBST, then serially dehydrated (10-15 minutes per wash) in 50%, 75%, and 100% ethanol diluted in distilled water. Tissue was stored at -20 °C in 100% ethanol until clearing and imaging. Tissue was cleared in BABB (1 part benzyl alcohol to 2 parts benzyl benzoate) for 2-12 hours at room temperature and mounted in BABB for imaging. For mounting, chambers were made on slides consisting of a thin layer of vacuum grease to create four walls followed by placement of a coverslip. Tissue was imaged on a Zeiss LSM 900 confocal microscope using the 10x objective.

For brain sections, brains were cryoprotected in 30% sucrose overnight. Tissue was cryosectioned (50 uM sections) and collected in a 12-well plate filled with PBS. Sections were stained free floating using a 12-well plate. Sections were washed 3x5 minutes each with 1x PBS containing 0.1% Triton X-100 (0.1% PBST), incubated with blocking solutions (0.1% PBST containing 5% normal goat serum (S-1000, Vector Labs) or normal donkey serum (005-000-121, Jackson Immuno) for 1-2 hours at room temperature, incubated with primary antibodies diluted in blocking solutions at 4 **°**C overnight. Subsequently, sections were washed 3x10 minutes each with 0.1% PBST, incubated with secondary antibodies diluted in blocking solutions at room temperature for 2 hours, and washed again 4x10 minutes each with 0.1% PBST (DAPI solution was included in the second wash at 1:5000 dilution). Sections were mounted onto glass slides and coverslipped in Vectashield (Vector Labs).

### Quantification of peripheral anatomy

For quantification of Meissner corpuscles in Ret cKO mice and littermate controls, cryosectioned and stained skin from the pads and fingertips were imaged with a widefield Zeiss LSM 700 microscope. Images of sections were analyzed in the Fiji distribution of Image J (www.fiji.sc). In each section, the number of Meissner corpuscles, visualized by S100 antibody staining, were counted. The length of the surface of the skin was traced and measured in ImageJ, and density was calculated by dividing the number of Meissner corpuscles by the length of the skin surface. Quantification was done blind to genotype.

For quantification of Pacinian corpuscles in Ret cKO mice, TrkB cKO mice, and littermate controls, images of stained and intact limbs were taken using a Zeiss LSM 900 confocal microscope using the 10x objective. The number of Pacinian corpuscles, visualized by S100 antibody staining, were counted. Quantification was done blind to genotype.

### DRG recordings

Animals were anesthetized with urethane (1.5 g/kg) or isoflurane (1.5-2%). A skin incision was made over the L3-L5 vertebrae and muscle surrounding the vertebrae was removed. The L3 and L5 vertebrae were fixed in place using custom-built clamps. The L4 DRG was exposed by removing overlaying bone. A borosilicate glass electrode (1 MΩ) was lowered into the DRG using a motorized manipulator (Scientifica) to search for units. Recordings were amplified using a MultiClamp 700B (Molecular Devices) under the 100x AC differential amplification mode with additional 20x gain. Signals were collected with a 0.1 kHz high-pass filter and 3 kHz Bessel filter and acquired using a Digidata 1550B (Molecular Devices) sampled at 20 kHz. For Pacinian corpuscle innervating Aβ-RA2-LTMRs, a search stimulus of 300 Hz vibration (40 mN) was applied to the ankle. Pacinians were identified by their large receptive fields, and their end organ location on the fibula was identified through electrical stimulation following measurements of their vibration tuning. For Meissner corpuscle innervating Aβ-RA1-LTMRs, the hindlimb digits were brushed as the electrode was descended. Aβ-RA1-LTMRs were identified by their small skin receptive fields, adapting responses to indentation, low mechanical thresholds (<10 mN), and cutaneous endings that could be excited by electrical stimulation of the skin. Vibrations (10-900 Hz) of varying intensity (1-50 mN) were delivered to the most sensitive area of the receptive field for each unit, collecting at least 4 trials per force for each frequency.

### Craniotomies for MEA recordings

Mice were injected with <u>sustained release</u> buprenorphine (0.1 mg/kg), anesthetized with isoflurane, and placed in a small animal stereotaxic frame (David Kopf Instruments). A titanium headplate was cemented to the mouse’s skull (C&B Metabond). A craniotomy was made centered over the right inferior colliculus (stereotaxic coordinates: 0.5 mm posterior, 1.8 mm lateral of lambda), or centered over the right VPL (stereotaxic coordinates: 1.7 mm posterior, 1.9 mm lateral of bregma), keeping the dura intact. Kwik-sil adhesive (World Precision Instruments) was used to cover the craniotomy until the day of recording (within 3 days of the craniotomy).

### Multielectrode array (MEA) recordings

For anesthetized recordings, the mouse was injected intraperitoneally with urethane (0.5-1.5 mg/g), placed on a heated platform, and head fixed such that the paws on the side contralateral to the craniotomy were accessible for mechanical stimulation. MEAs were inserted into the LCIC (H6b-32ch or H7b-32ch, Cambridge Neurotech) or VPL (H6b-32ch, H7b-32ch, or E4-64ch, Cambridge Neurotech) with a 3-axes micromanipulator (IVM mini, Scientifica). MEA signals were high-pass filtered at 200 Hz, amplified (RHD2132), and acquired at 20 kHz (RHD2000, Intantech) for offline processing.

For VPL recordings, hindlimb VPL was targeted using coordinates (1.6 mm posterior, 1.9 mm lateral to bregma), and the location was verified by brushing the hindlimb. The LCIC was targeted by positioning the MEA probe posterior to the transverse sinus and ∼1.6-1.8 mm lateral, with a 45-degree angle to insert the probe underneath the transverse sinus. In preliminary experiments, probe penetrations that contained only auditory responsive neurons were shown to be in the CNIC, unlike penetrations in the LCIC that contained both mechanical and auditory responses. Therefore, subsequent penetrations that contained only auditory responses were assumed to be in the CNIC and were not included in the analysis. For verification of recording location, after recording was complete, the probe was lifted from the brain, the probe was coated with DiI (Thermo Fisher) then re-inserted to the same coordinates, and the brain was collected and post-fixed overnight in 4% PFA post-recording. Recordings were performed and analyzed blind to genotype.

#### Force-frequency tuning experiments (anesthetized)

The receptive fields for neurons on the probe were determined by gently brushing the digits or pads using a fine paintbrush. The paw was fixed to the platform with Easy Mold Silicone Putty (Environmental Technologies). The mechanical stimulator (with a blunt 1 mm probe) was then targeted to the receptive fields identified by brush. In the case of LCIC neurons, which had large receptive fields that lacked a hotspot, the probe was targeted to the heel of the hindlimb. As above, for verification of recording location, after recording was complete, the probe was lifted from the brain, the probe was coated with DiI (Thermo Fisher) then re-inserted to the same coordinates, and the brain was collected and post-fixed overnight in 4% PFA post-recording. Thresholds for each vibratory frequency were defined as the force required to elicit 20% of a unit’s maximum firing during the sustained portion of the stimulus (200 ms window beginning 50 ms after stimulus onset).

#### VPL and LCIC mapping experiments (anesthetized)

For VPL mapping experiments (Figure S3) a 4 shanked probe (E4-64 ch, Cambridge Neurotech) was positioned with the shanks aligned to the antero-posterior axis. Mapping began medially (at ∼1.7 mm lateral) and moved 100 μm lateral for each successive penetration. For each probe penetration, the contralateral hindlimb, thigh, trunk, forelimb, and face were stroked with a paintbrush. The hindlimb was fixed to the platform with Easy Mold Silicone Putty and stimulation at 50 Hz and 500 Hz was delivered to the hindpaw with a 10 mN intensity using a mechanical stimulator with a wide blunt tip (3 mm diameter). Complementary LCIC experiments used a single shank probe (H6b or H7b, Cambridge Neurotech), with probe penetrations also separated by 100 μm intervals. Before each probe penetration the probe was coated with DiI and recording locations were verified post-experiment. Neurons were considered brush responsive to a body region if their change in firing exceeded 10 Hz during the brush period. Similarly, neurons were considered responsive to 500 Hz vibration if their change in firing exceeded 10 Hz during the sustained portion (200 ms period beginning 50 ms after stimulus onset) of the stimulus.

#### Awake LCIC recordings

For awake LCIC recordings, animals were head-fixed, and the LCIC was targeted as described above. Mechanical stimulation was targeted to the platform the animal stood on rather than the limb. The platform was made of thin (1/8”) acrylic. Trials where the animal was moving were excluded.

### Mechanical and auditory stimuli

For *in vivo* electrophysiology, vibrations were generated using a DC motor with a blunt 1-mm or 3-mm diameter probe, as previously described^15,27^. The motor was driven by a custom-built current supply controlled by Wavesurfer and a multifunction I/O board (USB-6343, National Instruments), and static forces calibrated using a fine scale. For awake recordings, the mechanical stimulator was covered with a sound dampening foam tip, and placed underneath the acrylic platform the animal stands on. The magnitude of vibration was measured using an accelerometer attached to the bottom of the platform and reported in units of gravitational force (g). Brush stimuli were manually delivered with a 4 mm diameter brush at a rate of one stroke per 2 seconds in the rostral to caudal direction.

Auditory stimuli were amplified by an SA1 stereo power amplifier (Tucker-Davis Technologies) and delivered through an MF1 multi-field magnetic speaker (Tucker-Davis Technologies), positioned 14 cm from the ear contralateral to recording site. Auditory stimuli were generated in MATLAB and delivered using WaveSurfer and a multifunction I/O board (USB-6343, National Instruments). Broadband (white) noise (0-50 kHz) was delivered in 100 ms pulses (5 ms rise-fall time) at 65 dB SPL. White noise, vibration, and combined trials were interleaved. For awake recordings, trials where the animal was moving were excluded.

### Spike detection and cluster assignment

Data was converted into contiguous binary files (https://github.com/peltonen/kwik-tools) and processed using JRClust (Janelia). Spikes were detected and assigned to clusters corresponding to individual or multiples units with JRClust (https://github.com/JaneliaSciComp/JRCLUST). These initial clusters were manually curated to remove artifacts that did not correspond to spike waveforms, and clusters were examined and split if their PCA decompositions were not monodispersed. This collection of clusters was then examined, and merge operations performed if consideration of the average spike waveform and spiking cross-correlation indicated that two clusters corresponded to the same neuron. Manual curation typically resulted in putative isolated single units with intra-cluster waveform correlations greater than 0.9, inter-cluster waveform correlations less than 0.1, and clusters that did not meet these criteria were not considered in the dataset. Spike event times for each cluster were exported and processed in Python.

### Anterograde tracing experiments

For anterograde tracing experiments (Figure S1), brainstem DCN neurons of *Slc17a6^Cre^* (Vglut2) mice were transduced with either an AAV encoding synaptophysin-GFP, AAV1-CAG-FLEX-Synaptophysin-GFP-WPRE (1.088E+14gc/ml), or synaptophysin-tDT, AAV1-CAG-FLEX-Synaptophysin-tdTomato-WPRE (1.1482E+14gc/ml). Viruses were produced and packaged at the Boston Children’s Hospital Viral Core Facility. Mice were injected with <u>sustained release</u> buprenorphine (0.1 mg/kg), anesthetized with isoflurane and placed in a small animal stereotaxic frame (David Kopf Instruments) with their neck bent at a 45 degree angle. The dorsal column nuclei were exposed between the base of the skull and C1 through retraction of the paraspinal muscles. A small nick in the overlying dural membrane was made to allow access for the injection pipette. A borosilicate glass pipette was lowered to a depth of 0.5 mm in either the gracile (0.1 mm lateral to obex) or cuneate (0.5mm anterior to obex, 0.1 mm lateral to fourth ventricle) nucleus. 50-100 nL of AAV virus with fast green was injected at a rate of 50 nL min−1 using a Microinject system (World Precision Instruments). Three weeks after injection, mice were perfused, and brain tissue was collected for cryosectioning and immunohistochemistry.

### Auditory brainstem responses (ABR) and distortion product otoacoustic emissions (DPOAE)

All measurements and analyses were conducted blind to the animal’s experimental condition. For anesthesia, mice were given an IP injection of ketamine (100 mg/kg body weight, BW), xylazine (10 mg/kg BW), and acepromazine (0.3 mg/kg BW) and placed on their side on a 37°C metal heating pad (ATC-2000, World Precision Instruments). Experiments were performed in a sound-attenuating, electrically shielded chamber (IAC Acoustics). After anesthesia induction, eye lubricant (GenTeal Tears, Alcon) was applied to prevent drying. A ketamine boost (0.3 – 0.5 mg/kg BW) was used to maintain anesthesia at the first sign of movement or elevation in the signal rejection rate (rejection threshold was 15 µV).

An acoustic system consisting of a microphone and dual earphone assembly coupled to a probe tube was used to deliver and measure acoustic stimuli. This custom system was built by the Engineering Core of the Eaton-Peabody Laboratories (EPL) at Massachusetts Eye and Ear (MEE, Boston). The probe end of the acoustic system was placed ∼1 mm over the center of the ear canal. Stimuli were digitally generated and then amplified by an SA1 stereo power amp (Tucker Davis Technologies) to drive the acoustic system. The probe tube microphone was amplified by a custom microphone amplifier (EPL at MEE) and sent to a 24-bit I/O board (PXI-4461) installed into a PXI chassis (PXIe-1073, National Instruments).

Bioelectrical signals were recorded using three subdermal electrodes: an active electrode at the vertex, a reference electrode near the pinna, and a ground electrode near the tail. Signals were passed through a custom pre-amplifier (EPL at MEE) before digitization. Acoustic stimuli consisted of clicks and 5 ms tone pips with a repetition rate of 30 s^-1^ and a 0.5 ms rise-fall time at the following frequencies (in kHz): 8, 16, 22.6, and 32. Stimuli were presented in alternating polarity to cancel out the cochlear microphonic potential. Cochlear Function Test Suite (CFTS) software (version 2.37.2; MEE, Boston) amplified (x104), filtered (0.3–3 kHz passband), and averaged 512 responses at each sound pressure level (SPL). ABR thresholds were manually designated blind to genotype using the ABR Peak Analysis Software from the EPL at MEE.

DPOAEs were recorded in response to primary tones f1 and f2 at a frequency ratio of f2/f1 = 1.2, and a SPL level of L1 = L2 + 10 dB. For individual DPOAE trials, f2 consisted of 5.6, 8, 11.3, 16, 22.6, 32, 42.5, or 64 kHz tones that were incremented in 10 dB steps from 10 to 70 dB SPL. The cubic distortion product 2f1-f2 was extracted by Fourier analysis of the ear-canal sound pressure after waveform and spectral averaging. DPOAE threshold was defined as the f1 level required to produce a DPOAE of 5 dB SPL, except at 64 kHz where the threshold was defined as 10 dB SPL due to a higher noise floor.

### Muscimol silencing

For muscimol silencing experiments, mice were injected with <u>sustained release</u> buprenorphine (0.1 mg/kg), anesthetized with isoflurane, and placed in a small animal stereotaxic frame (David Kopf Instruments). A small section of skin above the rear of the skull was removed with surgical scissors and the skull was dried. The skin was glued to the skull using Vetbond (3M healthcare), and a small amount of dental cement (C&B Metabond) was used to cover the exposed region of the skull, leaving the area where the craniotomy would be drilled clear. A small craniotomy was drilled approximately 0.5 mm posterior, 1.8 mm lateral to lambda. The craniotomy was covered with Kwik-sil adhesive.

Animals were given 3-4 days to recover. On the day of the behavioral experiment, animals were anesthetized with isoflurane and placed in a small animal stereotaxic frame. A borosilicate glass pipette was used to bilaterally inject into the inferior colliculus (0.5-0.8 mm posterior to lambda, 1.5-1.8 mm lateral, 0.5 mm deep) with TMR-X muscimol dissolved in ACSF (1.5 mM, Invitrogen M23400). 50 nL injections were made in two locations per side (100 nL total per side). Control mice received the same injection volume of sterile saline. Injections were performed at a rate of 50 nL min−1 using a Microinject system (World Precision Instruments) and the pipette was allowed to sit in the brain for 3 minutes post-injection. Animals were allowed to recover from isoflurane and habituate to the behavior room for ∼1 hour, and the behavioral assay (vibration, temperature, or texture) was run within 2 hours. Behavioral assays were run blind to experimental group (saline or muscimol). Post-behavior, animals were perfused. Muscimol spread was confirmed post-hoc following vibratome sectioning the brain and imaging.

### Behavior methods for Openfield, Vibration Platform and Temperature Preference assays

To reduce anxiety and improve handling, animals were enriched with Bio-Serv igloos, tunnels, or chew shacks at least one month prior to all behavior testing. One week prior to behavioral testing, animals were moved to the testing room and home cage habituated to the behavior room to acclimate to room lighting (warm white, 2700k), sound (ambient 45-50 dB), odor (cleaned with ECOS unscented dish soap and 70% EtOH), and temperature (73 °F, 30% humidity).

Vibration, temperature, and texture assays were conducted in 3-4 month old CD-1 animals (unless otherwise noted). Vibration assays in Ret cKO and littermate controls were performed at 8 weeks of age and were performed blind to genotype.

#### Vibration Platform and Sound Only Test

Two days prior to testing, animals were moved to the testing room and habituated to investigator handling. Animals were group habituated to the black matte test chamber (12 in L x 6 in W x 7 in H x 0.25 in), center divided with an arched doorway and allowed to freely explore for 5 minutes. During the three-day room/chamber habituation, animals were also habituated to the vibration motor sound only. The vibration platform was elevated 6 inches above the base. Two flat surface vibration motors, (PUI Audo, ASX08604-SW-R 4 ohm, BST) were suspended above the base, resting on spring mounted posts, one motor on each side of the chamber to act as the suspension base for the two square white matte acrylic floors, (5.5 in L x 5.5 in W x 0.125 in D) to rest on and provide vibration stimulus to each floor when activated. The acrylic floors were separated by a thin gap and lightly attached to the motor. An accelerometer was attached to each floor to ensure vibration floor displacement was constant for each test animal. The entire floor was covered with a thin stretch white matte cloth napkin (WypAll L30, 05812) to prevent the animals from investigating the gap in the floor and edges of the chamber walls as well as maintain a clean odor free test environment. The black matte acrylic test chamber rested on four pillars (6 inches above the floor) to ensure the chamber walls snugly surrounded the vibration floors without direct contact to prevent the chamber from vibrating when the motor was activated during testing. The room and test platform were lined with odorless 1-inch thick Bonded Logic Natural Fiber Acoustic Sound Absorbing panels (Model #60600-11212).

#### Vibration Test

The test chamber was modified on test day by removing the front and side wall to create a semi-open and closed end (L-shape) to increase animals’ awareness of the floor surface when exploring both sides of the test chamber. The center divider with open doorway was left in place. The semi-closed side of the chamber was the non-stimulus side and the semi-open side was the vibration stimulus side. A custom Bonsai script was used to trigger the vibration motor and camera. The three-minute vibration test sequence was as follows: the animal was placed in the chamber facing the semi-open side which encouraged the animal to walk through the doorway and explore the newly semi-closed side of the chamber.

Stage-1: Start, allow one minute baseline explore of test chamber,

Stage-2: 30 second vibration stimulus on the semi-open side,

Stage-3: 30 second interim period with no stimulus,

Stage-4: 30 second vibration stimulus on the semi-open side,

Stage-5: 30 second final period with no stimulus.

For the sound only test, the motor was disconnected from floor contact and the motor sound only was played for the 30 second stimulus during stage-2 and stage-4. Ambient room and chamber decibel level was 45-50 dB. During sound only and vibration stimulus periods, the background sound in the testing room and chamber was set to 70 dB at 500 Hz. A top video recording device was used to track distance traveled and time exploring the two chambers.

#### Temperature Preference Test

Two days prior to testing, animals were moved to the testing room and home cage habituated to the behavior. Animals were group habituated for 5 minutes to the black matte test chamber (12 in L x 6 in W x 7 in H x 0.25 in), center divided with an arched doorway, resting on two metal plate floors (5.5 in L x 5.5 in W ea) both set to neutral temperature 30 °C. On the third day following room habituation, each animal was tested on contrasting temperatures, 30 °C (warm) vs 18 °C (cold). Each animal was placed in the test chamber on the warm side facing the doorway to the cold side to encourage the animals to walk through the doorway and freely explore both temperatures for 5 minutes. A top video recording device was used to track distance traveled and time exploring the two chambers.

#### Texture Preference Assay

Two days prior to testing, animals were moved to the testing room and home cage habituated to the behavior. Animals were group habituated for 10 minutes to the black matte test chamber (12 in L x 6 in W x 7 in H x 0.25 in) with no divider. On the third day following room habituation, each animal was tested on contrasting floors, construction paper (Tru-Ray Heavyweight, Pacon) vs coarse 60-grit sandpaper (ceramic version, 26060PGP-4, 3M). There was no divider between the smooth and rough textured floors. Each animal was placed in the test chamber on the smooth side and animals were given 10 minutes to freely explore both chambers. A top video recording was used to track distance traveled and time exploring the two chambers.

### Quantification and statistical analysis

Statistical tests were conducted in Python 3.7.9 using the SciPy stats module or in GraphPad Prism. Normality of the data was tested using the Shapiro-Wilk test. Comparisons between two independent groups were performed using the unpaired t-test (in the case of two groups distributed normally, with no significant difference between variances), Welch’s t-test (in the case of two groups distributed normally, but with unequal variances), or Mann-Whitney test (non-parametric data). When correction for multiple comparisons was necessary, the Bonferroni correction was used. An adjusted p value < 0.05 was considered significant. Additional details on sample sizes and statistical tests for each experiment can be found in figure legends and the main text.

## Acknowledgements

We thank A. Handler, H. Kwak, K. Lezgiyeva, R. Martinez-Garcia, J. Peng, K. Ng, A. Emanuel, and A. Shuster for comments on the manuscript. We thank A. Emanuel for code used for the MEA data analysis. This work was supported by a HHMI Hannah Gray fellowship (GER), NEI P30 Core Grant for Vision Research #EY012196 (OM), NIH grants F31 NS097344 (ELH) and R35 5R35NS097344-05 (DDG), the Edward R. and Anne G. Lefler Center for Neurodegenerative Disorders (DDG), and the Hock E. Tan and Lisa Yang Center for Autism Research (DDG). DDG is an investigator of the Howard Hughes Medical Institute. This article is subject to HHMI’s Open Access to Publications policy. HHMI lab heads have previously granted a nonexclusive CC BY 4.0 license to the public and a sublicensable license to HHMI in their research articles. Pursuant to those licenses, the author-accepted manuscript of this article can be made freely available under a CC BY 4.0 license immediately upon publication.

## Author Contributions

ELH and DDG conceived the study. ELH did the MEA recordings in both anesthetized and awake conditions, with help from JT and MD. JT and ZKS performed primary sensory neuron recordings. ELH, AH, and MD performed anatomy and immunohistochemistry experiments. MMD, OM, and ELH developed the vibration behavioral assay. ELH did the saline and muscimol IC injections, and MMD performed behavioral experiments for vibration, temperature, and texture preference assays, with help from ELH. GER performed auditory brainstem response (ABR) measurements with guidance from LVG. KTB and DPC generated and provided the *GJB2^flox/flox^ Sox10^Cre^* mice. ELH and DDG wrote the manuscript with input from all authors.

